# Inhibition stabilization and paradoxical effects in recurrent neural networks with short-term plasticity

**DOI:** 10.1101/2022.12.19.520986

**Authors:** Yue Kris Wu, Julijana Gjorgjieva

## Abstract

Inhibition stabilization is considered a ubiquitous property of cortical networks, whereby inhibition controls network activity in the presence of strong recurrent excitation. In networks with fixed connectivity, an identifying characteristic of inhibition stabilization is that increasing (decreasing) excitatory input to the inhibitory population leads to a decrease (increase) in inhibitory firing, known as the paradoxical effect. However, population responses to stimulation are highly nonlinear, and drastic changes in synaptic strengths induced by short-term plasticity (STP) can occur on the timescale of perception. How neuronal nonlinearities and STP affect inhibition stabilization and the paradoxical effect is unclear. Using analytical calculations, we demonstrate that in networks with STP the paradoxical effect implies inhibition stabilization, but inhibition stabilization does not imply the paradoxical effect. Interestingly, networks with neuronal nonlinearities and STP can transition non-monotonically between inhibition-stabilization and non-inhibition-stabilization, and between paradoxically- and non-paradoxically-responding regimes with increasing excitatory activity. Furthermore, we generalize our results to more complex scenarios including networks with multiple interneuron subtypes and any monotonically increasing neuronal nonlinearities. In summary, our work reveals the relationship between inhibition stabilization and the paradoxical effect in the presence of neuronal nonlinearity and STP, yielding several testable predictions.

## I. INTRODUCTION

Cortical networks are typically characterized by inhibition stabilization, where inhibition is needed to keep network activity levels in biologically realistic ranges despite the presence of strong recurrent excitation [1]. Networks operating in the inhibition-stabilized regime are capable of performing various computations, including input amplification, response normalization, and network multistability [2–6]. In networks with fixed connectivity, a hallmark of inhibition stabilization is the paradoxical effect: an increase or a decrease of excitatory input to the inhibitory population respectively decreases or increases the inhibitory firing [7]. Over the past decade, much effort has been made to identify the operating regime of cortical networks based on the paradoxical effect [1, 8, 9].

Yet, various aspects ranging from the network to the synaptic level can considerably affect network dynamics and the operating regime. First, if individual neurons in the network receive large excitatory and inhibitory currents which precisely cancel each other, the network operates in a balanced state characterized by a linear population response [10– 12]. Recent work has argued that neuronal inputoutput functions are better characterized by supralinear functions, and networks with this type of non-linearity can exhibit various nonlinear phenomena as observed in biology [13–15]. Second, synapses in the brain are highly dynamic as a result of different short-term plasticity (STP) mechanisms, operating on a timescale of milliseconds to seconds [16, 17]. Upon presynaptic stimulation, postsynaptic responses can either get depressed subject to short-term depression (STD) or facilitated subject to short-term facilitation (STF). While short-term synaptic dynamics are widely observed in biological circuits, it is unclear how they interact with the neuronal nonlinearity to jointly determine the network operating regime. Here we ask how the neuronal nonlinearity and STP affect inhibition stabilization and the paradoxical effect.

To address this question, we determine the conditions for inhibition stabilization and the paradoxical effect in networks of excitatory and inhibitory neurons in the presence of STP with linear and supralinear population response functions. We find that irrespective of the neuronal nonlinearity, in networks with excitatory-to-excitatory (E-to-E) STD, inhibition stabilization does not necessarily imply the paradoxical effect, but the paradoxical effect implies inhibition stabilization. In contrast, in networks with static connectivity or networks with other STP mechanisms, inhibition stabilization and the paradoxical effect imply each other. Interestingly, neuronal nonlinearities and STP endow the network with unconventional behaviors. More specifically, in the presence of a neuronal nonlinearity and E-to-E STD, monotonically increasing excitatory activity can lead to non-monotonic transitions between non-inhibition-stabilization and inhibition-stabilization, as well as between non-paradoxically-responding and paradoxically-responding regimes. Furthermore, we generalize our results to more complex scenarios including networks with multiple interneuron subtypes and any monotonically increasing neuronal nonlinearities. In conclusion, our work reveals the impact of neuronal nonlinearities and STP on inhibition stabilization and the paradoxical effect, and makes several predictions for future experiments.

## II. RESULTS

To understand the relationship between inhibition stabilization and the paradoxical effect in recurrent neural networks with STP, we studied rate-based population models consisting of an excitatory (E) and an inhibitory (I) population. The dynamics of the network are given by:

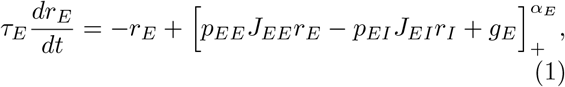

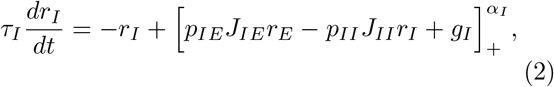

where *r*_*E*_ and *r*_*I*_ denote the firing rates of the excitatory and inhibitory population, *τ*_*E*_ and *τ*_*I*_ are the corresponding time constants, *J*_*AB*_ represents the synaptic strength from population *B* to population

*A*, where *A, B ∈ {E, I}, g*_*E*_ and *g*_*I*_ are the external inputs to the respective populations, *α*_*E*_ and *α*_*I*_ are the exponents of the respective input-output functions. Finally, *p*_*AB*_ represents the short-term plasticity variable from population *B* to population *A*. We implemented short-term plasticity mechanisms based on the Tsodyks and Markram model [16]. For STD, we replaced *p*_*AB*_ by *x*_*AB*_ and described the STD dynamics as follows:

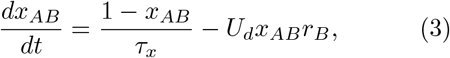

where *x*_*AB*_ is a short-term depression variable that is limited to the interval (0,1] for the synaptic connection from population B to population A. Biophysically, the short-term depression variable *x* represents the fraction of vesicles available for release, *τ*_*x*_ is the time constant of STD, and *U*_*d*_ is the depression factor controlling the degree of depression induced by the presynaptic activity.

For STF, we replaced *p*_*AB*_ by *u*_*AB*_ and expressed the STF dynamics as follows:

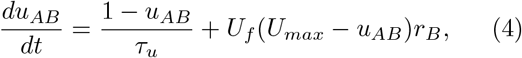

where *u*_*AB*_ is a short-term facilitation variable that is constrained to the interval [1, *U*_*max*_) for the synaptic connection from population B to population A. Biophysically, the short-term facilitation variable *u* represents the ability of releasing neurotransmitter, *τ*_*u*_ is the time constant of STF, *U*_*f*_ is the facilitation factor controlling the degree of facilitation induced by the presynaptic activity, and *U*_*max*_ is the maximal facilitation value.

To investigate the impact of neuronal nonlinearities on inhibition stabilization and the paradoxical effect, we considered both threshold-linear networks (*α*_*E*_ = *α*_*I*_ = 1) as well as supralinear networks (*α*_*E*_ = *α*_*I*_ *>* 1). In the regime of positive *r*_*E*_ and *r*_*I*_, threshold-linear networks behave as linear networks. In the following, we thus call them linear networks. Furthermore, while we keep our analysis for supralinear networks in a general form, we use *α*_*E*_ = *α*_*I*_ = 2 for the numerical simulations. Note that the neuronal nonlinearity in our study refers to the nonlinearity of population-averaged responses to input when no STP mechanisms are taken into account, which is fully determined by *α*_*E*_ and *α*_*I*_ .

In addition, for the sake of analytical tractability, we included one STP mechanism at a time. To investigate how inhibition stabilization is affected by the neuronal nonlinearity and STP, we computed the real part of the leading eigenvalue of the Jacobian matrix of the excitatory-to-excitatory subnetwork incorporating STP, and refer to it as the ‘Inhibition Stabilization index’ (IS index) (Supplemental Methods; Supplemental Table 1). A positive (negative) IS index implies that the network is in the IS (non-IS) regime. To reveal how inhibition stabilization changes with network activity and network connectivity, we investigated how the IS index changes with the excitatory activity *r*_*E*_ and the excitatory to excitatory connection strength *J*_*EE*_. These two quantities, *r*_*E*_ and *J*_*EE*_, are directly involved in the definition of the IS index (Supplemental Table 1).

### A. INHIBITION STABILIZATION IN RECURRENT NEURAL NETWORKS WITH SHORT-TERM DEPRESSION AT E-TO-E SYNAPSES

We first examined inhibition stabilization for networks with E-to-E STD, evaluated at the fixed point of the system (Fig. 1a). The distinction between non-IS and IS is reflected in network responses to perturbations induced by injecting additional excitatory currents into excitatory neurons while inhibition is fixed. Networks initially in the non-IS regime return back to their initial activity level after a small transient perturbation to the excitatory activity when inhibition is fixed, whereas networks initially in the IS regime deviate from their initial activity (Fig. S1). For linear networks with E-to-E STD, if *J*_*EE*_ is less than 1, the network is always in the non-IS regime regardless of *r*_*E*_ (Fig. 1b). If *J*_*EE*_ is greater than 1, the network transitions from IS to non-IS with increasing *r*_*E*_ (Fig. 1b). In contrast, supralinear networks with E-to-E STD manifest different behaviors. When *J*_*EE*_ is large, the network first transitions from non-IS to IS, and then back to non-IS with increasing *r*_*E*_ (Fig. 1c, Fig. S1). When *J*_*EE*_ is small, the supralinear network stays in the non-IS regime for all values of *r*_*E*_ (Fig. 1c).

**FIG. 1.**
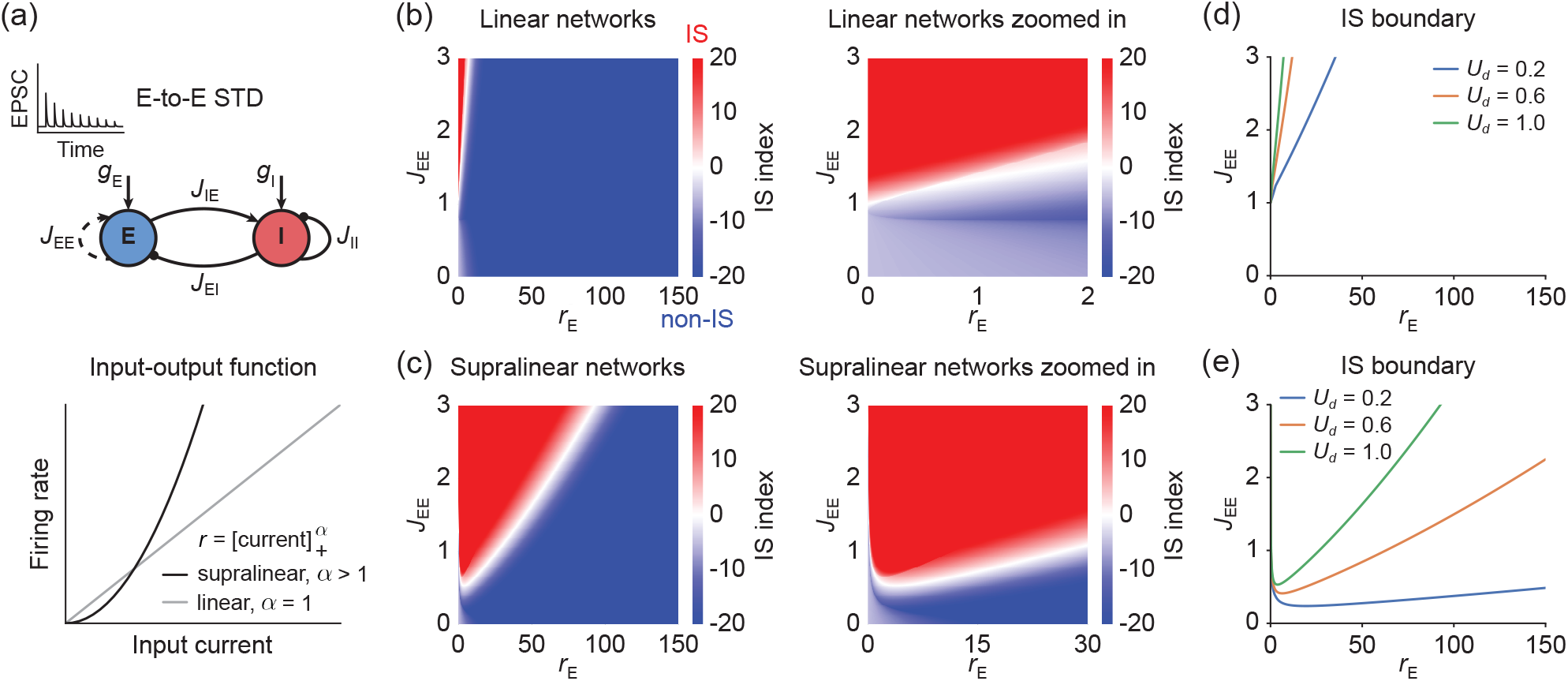
Inhibition stabilization in recurrent neural networks with E-to-E short-term depression. (a) Top: Schematic of the recurrent network model consisting of an excitatory (blue) and an inhibitory population (red) with E-to-E STD. Bottom: Different nonlinearities of population-averaged responses to input. Linear (gray) and supralinear (black) input-output function given by a rectified power law with exponent *α* = 1 and *α* = 2 (cf. Eq. (1)-(2)), respectively. (b) Left: IS index as a function of excitatory firing rate *r*_*E*_ and excitatory-to-excitatory connection strength *J*_*EE*_ for linear networks with E-to-E STD. The non-IS and IS regime are marked in blue and red, respectively. Right: zoomed-in version of (b) left. Here, the depression factor *U*_*d*_ is 1.0 (Supplemental Methods), the firing rate *r*_*E*_ is plotted in a biologically realistic range from 0 to 150 Hz. (c) Same as (b) but for supralinear networks with E-to-E STD. Here, the depression factor *U*_*d*_ is 1.0 (Supplemental Methods). (d) IS boundary for linear networks with E-to-E STD, defined as the corresponding excitatory-to-excitatory connection strength 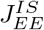 for different *r*_*E*_ at which the IS index is 0. Different colors represent the IS boundary for different values of depression factor *U*_*d*_. (e) Same as (d) but for supralinear networks with E-to-E STD. Here, *τ*_*x*_ is 200 ms in (b-e).

To better understand the transition between non-IS and IS in the presence of neuronal nonlinearities and E-to-E STD, we investigated how the boundary between non-IS and IS, defined as ‘IS boundary’, changes with *r*_*E*_ (Supplemental Methods; Fig. S2). Mathematically, the IS boundary is determined by the recurrent excitatory-to-excitatory connection strength for different *r*_*E*_ at which the IS index is 0, denoted by 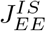. By computing the derivative of 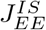 with respect to *r*_*E*_ (Supplemental Methods), we found that the derivative is always positive for linear networks with E-to-E STD, suggesting that the IS boundary increases with increasing *r*_*E*_ (Fig. 1d). Therefore, for networks with large *J*_*EE*_, as *r*_*E*_ increases, only the transition from IS to non-IS is possible (Fig. 1b, d). In contrast, for supralinear networks with E-to-E STD, the derivative changes from negative to positive with increasing *r*_*E*_ (Supplemental Methods), implying that the IS boundary first decreases and then increases as *r*_*E*_ increases (Fig. 1e). As a result, networks can undergo non-monotonic transitions between non-IS and IS with increasing *r*_*E*_. More specifically, networks can switch from non-IS to IS, and then back to non-IS with increasing *r*_*E*_ (Fig. 1c, e; Fig. S1). The non-monotonic transitions arise from the competition between the increasing neuronal gain due to the supralinear neuronal input-output function and the decreasing synaptic strength due to STD. Despite high firing rates in the large *r*_*E*_ limit, E-to-E synaptic strengths are significantly depressed and STD effectively decouples excitatory neurons, rendering the network non-inhibition stabilized. Furthermore, the turning point of the IS boundary appears when *U*_*d*_*x*_*EE*_*r*_*E*_ is of order 1 (Supplemental Methods). Increasing the depression factor *U*_*d*_ or the STD time constant *τ*_*x*_ shifts the turning point to the upper left in the (*r*_*E*_, *J*_*EE*_) coordinate system (Fig. S3; Supplemental Methods). We also found that these non-monotonic transitions cannot be observed in networks with static connectivity (Fig. S4; Supplemental Methods). Taken together, our results suggest that E-to-E STD can nontrivially affect the inhibition stabilization property, especially in the presence of neuronal nonlinearities.

### B. INHIBITION STABILIZATION IN RECURRENT NEURAL NETWORKS WITH SHORT-TERM FACILITATION AT E-TO-E SYNAPSES

To determine if the observed effects are specific to the type of STP at E-to-E synapses, we next examined networks with E-to-E STF (Fig. 2a). Unlike the scenario with STD, for both linear networks or supralinear networks, only a monotonic transition from non-IS to IS is possible with increasing *r*_*E*_ in the presence of E-to-E STF (Fig. 2b, c). In contrast to supralinear networks, linear networks with *J*_*EE*_ larger than 1 are always in the IS regime regardless of *r*_*E*_. In both cases, the parameter regime of *J*_*EE*_ and *r*_*E*_ which supports IS is much larger than in the corresponding network with STD (Fig. 2). Furthermore, independent of the neuronal nonlinearity, the derivative of 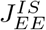 with respect to *r*_*E*_ is always negative (Supplemental Methods), indicating that the IS boundary decreases as *r*_*E*_ increases (Fig. 2d, e). These results indicate that E-to-E STF exerts a more intuitive influence on the inhibition stabilization property than E-to-E STD even in the presence of neuronal nonlinearities.

**FIG. 2.**
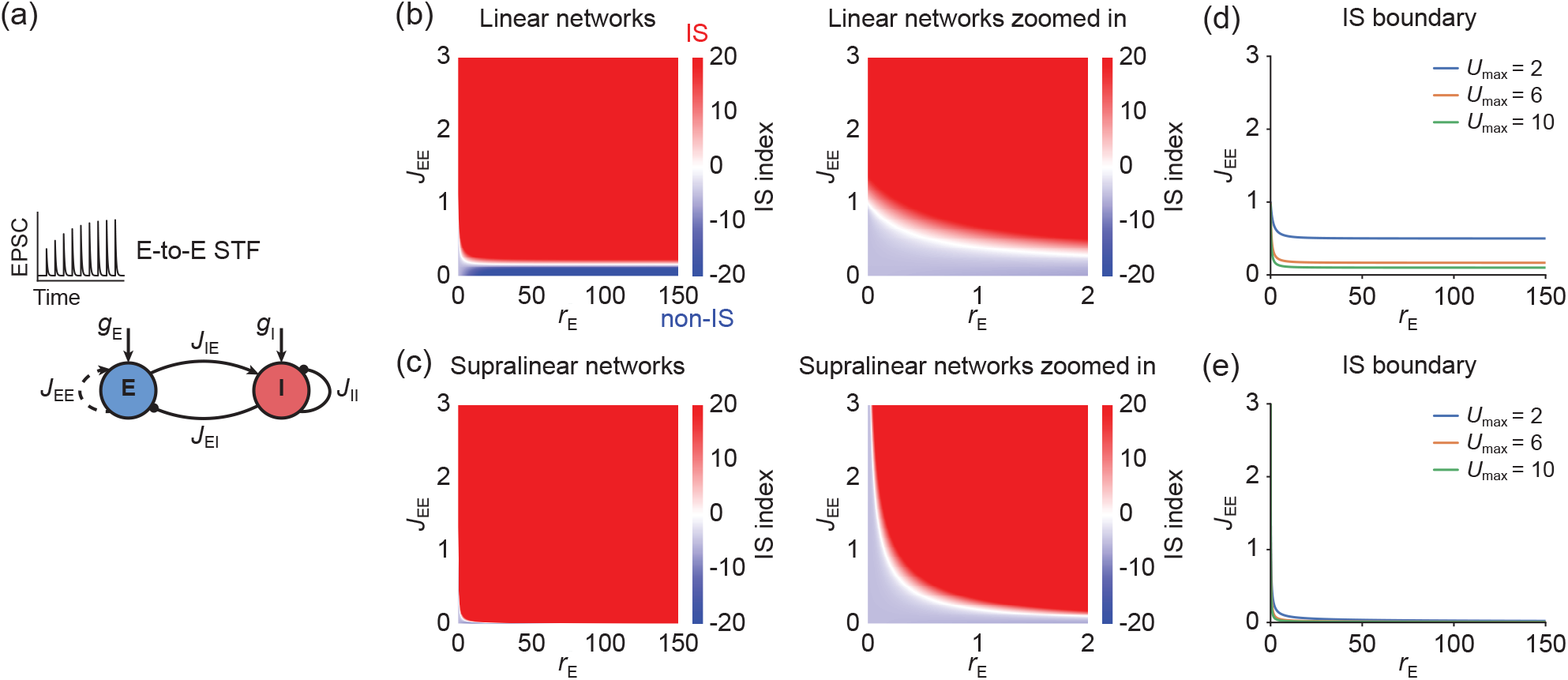
Inhibition stabilization in recurrent neural networks with E-to-E short-term facilitation. (a) Schematic of the recurrent network model consisting of an excitatory (blue) and an inhibitory population (red) with E-to-E STF. (b) Left: IS index as a function of excitatory firing rate *r*_*E*_ and excitatory-to-excitatory connection strength *J*_*EE*_ for linear networks with E-to-E STF. The non-IS and IS regime are marked in blue and red, respectively. Right: zoomed-in version of (b) left. Here, the facilitation factor *U*_*f*_ is 1.0, and the maximal facilitation value *U*_*max*_ is 6.0 (Supplemental Methods). (c) Same as (b) but for supralinear networks with E-to-E STF. Here, the facilitation factor *U*_*f*_ is 1.0, and *U*_*max*_ is 6.0 (Supplemental Methods). (d) IS boundary for linear networks with E-to-E STF, defined as the corresponding excitatory-to-excitatory connection strength 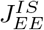 for different *r*_*E*_ at which the IS index is 0. Different colors represent the IS boundary for different values of *U*_*max*_. (e) Same as (d) but for supralinear networks with E-to-E STF. Here, *τ*_*u*_ is 200 ms in (b-e).

### C. INHIBITION STABILIZATION IN RECURRENT NEURAL NETWORKS WITH SHORT-TERM PLASTICITY AT OTHER SYNAPSES

Finally, we performed the same analyses for networks with different types of STP at all synapses other than E-to-E, including E-to-I STD/STF, I-to-E STD/STF, and I-to-I STD/STF, respectively (Fig. 3a; Supplemental Methods). Including these STP mechanisms does not change the IS condition relative to networks with fixed connectivity (Supplemental Methods). For networks with a linear input-output function, the IS boundary does not change with *r*_*E*_ (Fig. 3b, d; Supplemental Methods), and *J*_*EE*_ completely determines whether the network is non-IS or IS. In contrast, for networks with a supralinear input-output function, the derivative of 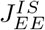 with respect to *r*_*E*_ is always negative, suggesting that the IS boundary decreases with increasing *r*_*E*_ (Fig. 3c, e; Supplemental Methods). Therefore, the transition from non-IS to IS with increasing *r*_*E*_ in static supralinear networks or supralinear networks with STP at all synapses other than E-to-E can only happen for large *J*_*EE*_ (Fig. 3c, e; Fig. S4). No transition between non-IS and IS can occur in the biological realistic firing regime from 0 to 150 Hz for small *J*_*EE*_ (Fig. 3c, e).

**FIG. 3.**
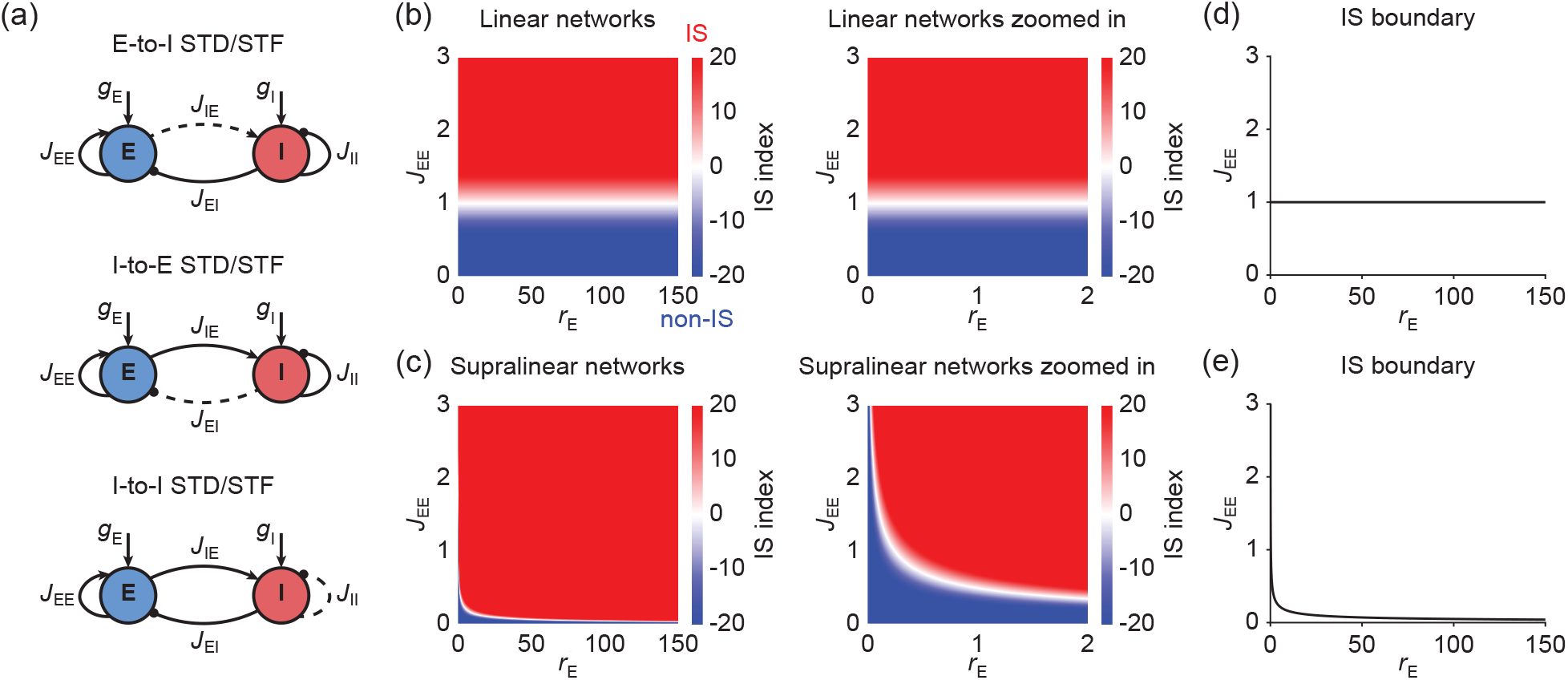
Inhibition stabilization in recurrent neural networks with E-to-I short-term depression/facilitation, I-to-E short-term depression/facilitation, or I-to-I short-term depression/facilitation. (a) Schematic of the recurrent network model consisting of an excitatory (blue) and an inhibitory population (red) with E-to-I STD/STF (top), with I-to-E STD/STF (middle), or with I-to-I STD/STF (bottom). (b) Left: IS index as a function of excitatory firing rate *r*_*E*_ and excitatory-to-excitatory connection strength *J*_*EE*_ for linear networks with E-to-I STD/STF, with I-to-E STD/STF, or with I-to-I STD/STF. The non-IS and IS regime are marked in blue and red, respectively. Right: zoomed-in version of (b) left. (c) Same as (b) but for supralinear networks with E-to-I STD/STF, with I-to-E STD/STF, or with I-to-I STD/STF. (d) IS boundary for linear networks with E-to-I STD/STF, with I-to-E STD/STF, or with I-to-I STD/STF, defined as the corresponding excitatory-to-excitatory connection strength 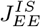 for different *r*_*E*_ at which the IS index is 0. (e) Same as (d) but for supralinear networks with E-to-I STD/STF, with I-to-E STD/STF, or with I-to-I STD/STF.

In summary, by considering all possible STP mechanisms, our results demonstrate a nontrivial influence of the neuronal nonlinearity and STP on inhibition stabilization. Specifically, we revealed how inhibition stabilization changes with excitatory activity and network connectivity when considering neuronal nonlinearities and STP.

### D. PARADOXICAL EFFECTS IN RECURRENT NEURAL NETWORKS WITH SHORT-TERM PLASTICITY

Previous theoretical studies have suggested that in excitatory and inhibitory networks with static connectivity one identifying characteristic of inhibition stabilization is that injecting excitatory (inhibitory) current into inhibitory neurons decreases (increases) inhibitory firing rates, known as the paradoxical effect [4, 7]. Here, we sought to identify the conditions under which a paradoxical effect can arise in recurrent neural networks with STP. We assumed that the system is stable locally around the fixed point, in other words, a small transient perturbation to the system will not lead to deviation from the initial fixed point over time. Furthermore, the perturbation used to probe the paradoxical effect (e.g. the excitatory current injected to the inhibitory population) is small enough that it will not lead to a transition between non-IS and IS. To determine the conditions for the presence of the paradoxical effect under these assumptions, we considered the phase plane of the excitatory (abscissa) and inhibitory (ordinate) firing rate dynamics. The first condition involves a larger slope of the inhibitory nullcline than of the excitatory nullcline locally around the fixed point in the phase plane, while the second condition involves a positive slope of the excitatory nullcline around the fixed point (7; Fig. S5; Supplemental Methods). We compared the above two conditions for the presence of the paradoxical effect with the conditions to be in the IS regime. We found that irrespective of the shape of the neuronal nonlinearity, in networks with E-to-E STD, the paradoxical effect implies inhibition stabilization, whereas inhibition stabilization does not imply the paradoxical effect (Fig. 4a; Supplemental Methods). In contrast, for networks with E-to-E STF, E-to-I STD/STF, I-to-I STD/STF, and I-to-I STD/STF, inhibition stabilization and the paradoxical effect imply each other (Fig. 4a; Supplemental Methods). To highlight the difference between inhibition stabilization and the paradoxical effect in networks with E-to-E STD (Fig. 4b), we plotted the paradoxical effect boundary that separates the paradoxically-responding and the non-paradoxically-responding regime together with the IS boundary for both linear networks and supralinear networks with E-to-E STD (Fig. 4c-f). In the two dimensional *r*_*E*_-*J*_*EE*_ plane, the parameter regime for the paradoxical effect is much narrower than the IS regime, suggesting that there is a large parameter space, in which inhibition-stabilized networks do not exhibit the paradoxical effect (Fig. 4c-f). It is noteworthy that in the presence of E-to-E STD, networks in which dynamic STD regulation is required to ensure stability, as studied in [18, 19], are a subset of inhibition stabilized networks which do not exhibit paradoxical effects (Supplemental Methods).

**FIG. 4.**
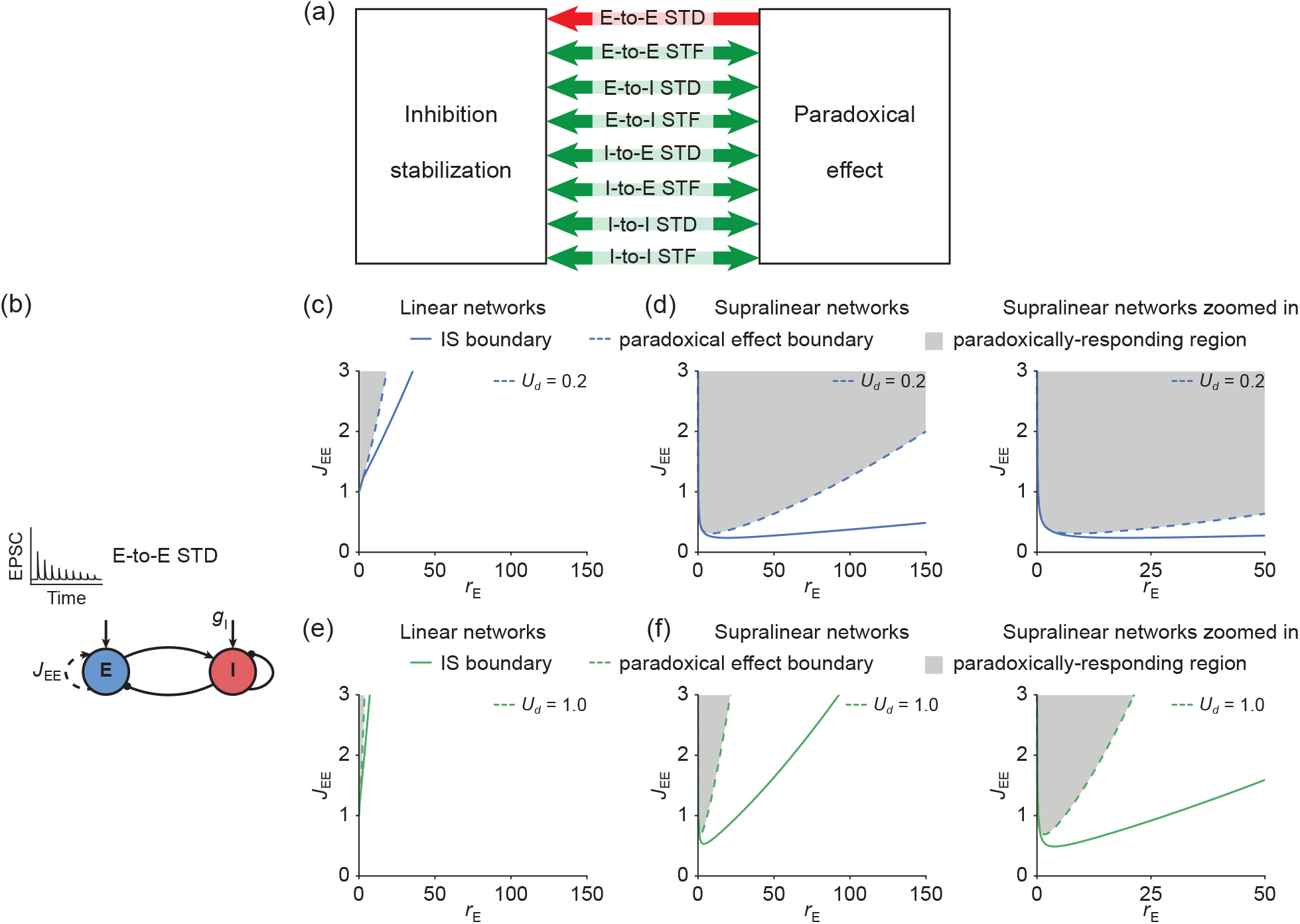
Relationship between inhibition stabilization and the paradoxical effect in recurrent neural networks with short-term plasticity. (a) In networks with E-to-E STD, the paradoxical effect implies inhibition stabilization, whereas inhibition stabilization does not imply the paradoxical effect indicated by the unidirectional red arrow. In contrast, for networks with E-to-E STF, E-to-I STD/STF, I-to-I STD/STF, and I-to-I STD/STF, inhibition stabilization and the paradoxical effect imply each other indicated by the bidirectional green arrows. (b) Schematic of the recurrent network model consisting of an excitatory (blue) and an inhibitory population (red) with E-to-E STD. (c) An example of the paradoxical effect boundary (dashed line) and the inhibition stabilization boundary (solid line) as a function of excitatory firing rate *r*_*E*_ for linear networks with E-to-E STD. The paradoxically-responding region is marked in gray. Here, the depression factor *U*_*d*_ is 0.2 (Supplemental Methods). (d) Same as (c) but for supralinear networks with E-to-E STD. Right: zoomed-in version of (d) left. (e-f) Same as (c-d) but the depression factor *U*_*d*_ 1.0. Here, *τ*_*x*_ is 200 ms in (c-f).

Furthermore, by analyzing how the paradoxical effect boundary changes with *r*_*E*_, we found that it exhibits a similar trend to the IS boundary (Supplemental Methods). In particular, the paradoxical effect boundary of supralinear networks with E-to-E STD is also a non-monotonic function of *r*_*E*_. Therefore, in this case, networks can undergo non-monotonic transitions between the paradoxically-responding regime and non-paradoxically-responding regime with monotonically changing excitatory activity *r*_*E*_.

### E. GENERALIZATION TO NETWORKS WITH A MIXTURE OF STP MECHANISMS AND MULTIPLE INTERNEURON SUBTYPES

So far, we only considered individual STP mechanisms one at a time. Here, we generalized our findings to four more complex and biologically realistic scenarios. First, our results can be generalized to networks with a mixture of STP mechanisms at other types of synapses except E-to-E synapses where we assume only STD or STF (Supplemental Methods). More specifically, the conditions and results derived for networks with either E-to-E STD or STF alone are the same for networks with either E-to-E STD orSTF and a mixture of STP mechanisms at other types of synapses.

Second, by incorporating both STD and STF at E-to-E synapses, we found that the paradoxical effect implies inhibition stabilization, whereas inhibition stabilization does not necessarily imply the paradoxical effect (Supplemental Methods).

Third, inhibitory neurons in the cortex are highly diverse in terms of anatomy, electrophysiology and function [20–22]. Recent studies have investigated the relationship between inhibition stabilization and the paradoxical effect in networks with multiple interneuron subtypes in the absence of STP [23–27]. Yet, synapses between different interneuron subtypes exhibit considerable short-term dynamics [28]. We then extended the analysis to networks with multiple interneuron subtypes. Theoretical studies have shown that in networks with static connectivity, if the excitatory subnetwork is non-IS (IS), then the sign of the change in the firing rate of the excitatory population and of the change in the total inhibitory current to the excitatory population are opposite (the same) [23]. We found that in the presence of E-to-E STD, if the network is IS, the sign of the change in the firing rate of the excitatory population and of the change in the total inhibitory current to the excitatory population are the same (Supplemental Methods). However, the same sign of the change in the firing rate of the excitatory population and of the change in the total inhibitory current to the excitatory population does not imply that the network is IS (Supplemental Methods). In the presence of E-to-E STF and STP at other types of synapses, inhibition stabilization and the same sign change in the firing rate of the excitatory population and of the change in the total inhibitory current to the excitatory population imply each other (Supplemental Methods).

Last, the derived relationship between inhibition stabilization and the paradoxical effect is independent from the power of the power-law input-output function. Therefore, the derived relationship holds for any monotonically increasing neuronal nonlinearities including sublinear (Fig. S6, Supplemental Methods), despite the fact that sublinear networks have different IS transitions from supralinear networks (Fig. S7, Supplemental Methods). Taken together, our results indicate that the relationship between inhibition stabilization and paradoxical effect in networks with STP becomes non-trivial in the presence of short-term plasticity.

## III. DISCUSSION

In this study, we combined analytical and numerical methods to reveal the effects of neuronal nonlinearities and STP on inhibition stabilization, the paradoxical effect, and the relationship between them. Including STD at E-to-E synapses, in contrast to other types of STP and other synapse types, generates the most surprising results. Under these conditions, the paradoxical effect implies inhibition stabilization, whereas inhibition stabilization does not imply the paradoxical effect. For networks with other STP mechanisms and networks with static connectivity, inhibition stabilization and the paradoxical effect imply each other. Additionally, in the presence of a neuronal nonlinearity and E-to-E STD, a non-monotonic transition between different regimes can occur when excitatory activity changes monotonically.

Network models applied to neural circuit development have previously investigated inhibition stabilization in the presence of STP [29]. Recent studies have examined inhibition stabilization and the paradoxical effect in threshold-linear networks with E-to-E STP to demonstrate that inhibition stabilization can be probed by the paradoxical effect [1]. Recent work has conducted similar analysis for supralinear networks with short-term plasticity on specific synapses [30]. Here, we systematically analyzed networks with all forms of STP mechanisms, for both linear networks and supralinear networks. By mathematically defining the IS boundary and the paradoxical effect boundary, we further demonstrated how network activity and connectivity affect the inhibition stabilization property and the paradoxical effect. Importantly, we generalized our results to several more complex scenarios including networks with a mixture of STP mechanisms, networks with both E-to-E STD and STF, networks with multiple interneuron subtypes, and any monotonically increasing neuronal nonlinearities.

In this work, we assumed that the network activity has reached a fixed point, and we did not consider scenarios like multistability or oscillations that could arise from neuronal nonlinearities or STP [15, 31, 32]. While multistability and oscillations have been observed in the brain [33, 34], the single stable fixed point assumed in our study is believed to be a reasonable approximation of awake sensory cortex [35].

Our model makes several predictions that can be tested in optogenetic experiments. Across cortical layers and across brain regions, synaptic strengths can differ by an order of magnitude [36]. Furthermore, the degrees of balance between excitation and inhibition may also vary [37, 38], resulting different neuronal nonlinearities [11, 13, 38]. Therefore, different behaviors predicted by our analysis may be observable in different neural circuits. For example, in the presence of E-to-E STD, our model shows that networks with weak excitatory to excitatory connection strength *J*_*EE*_ are always nonIS in the biologically realistic activity regime and therefore exhibit no paradoxical effects. In contrast, with E-to-E STD and strong *J*_*EE*_, network models with a linear population-averaged response function can undergo the transition from IS to nonIS with increasing excitatory activity *r*_*E*_. Different from linear networks, our model predicts that nonmonotonic transitions between non-IS and IS can be found in supralinear networks. More specifically, supralinear networks can switch from non-IS to IS, and then from IS to non-IS with increasing *r*_*E*_. Although inhibition stabilization does not imply the paradoxical effect in the presence of E-to-E STD, the transition between paradoxically-responding and non-paradoxically-responding regime is also nonmonotonic with increasing *r*_*E*_ in supralinear networks, whereas in linear networks, only transitions from the paradoxically-responding to the nonparadoxically-responding regime with increasing *r*_*E*_ is possible. Therefore, depending on the excitation-inhibition balance, and the specific short term plasticity mechanisms operating in different brain regions, our work proposes that the circuits will exhibit different properties when interrogated with common experimental techniques.

Second, in the presence of STF on E-to-E synapses or STP on other synapses, our results demonstrate that inhibition stabilization and the paradoxical effect imply each other. Linear network models with *J*_*EE*_ larger than 1 that have E-to-E STF are always IS and thus exhibit the paradoxical effect. In linear network models with STP on other synapses, activity does not affect inhibition stabilization and the paradoxical effect. In contrast, regardless of the strength of *J*_*EE*_, supralinear networks with E-to-E STF or STP on other synapses can switch from non-IS to IS with increasing *r*_*E*_. This transition from non-IS and IS can be directly tested experimentally by probing the paradoxical effect, because of the equivalence of inhibition stabilization and the paradoxical effect found in network models with E-to-E STF or STP on other synapses. Last, our analysis shows that in most cases substantially altering either *J*_*EE*_ or *r*_*E*_ can switch the network operating regime. Multiple factors can modify *J*_*EE*_ and *r*_*E*_ experimentally. On a short timescale, the strength of sensory stimulation, and behavioral states like arousal [39], attention [40] and locomotion [41] can dramatically change activity levels *r*_*E*_. Regime switching may therefore be experimentally observable across different stimulation conditions and different behavioral states. On a long timescale, *J*_*EE*_ or *r*_*E*_ can be modified by long term plasticity mechanisms [42, 43]. In this case, regime switching could be experimentally detectable across different developmental stages.

Taken together, our theoretical framework provides a systematic analysis of how short-term synaptic plasticity and response nonlinearities interact to determine the network operating regime, revealing unexpected relationships and their signatures as a guide for future experimental studies.

## ACKNOWLEDGMENTS

We thank Elizabeth Herbert, Felix Waitzmann, Leonidas M. A. Richter, and Dylan Festa for comments on the manuscript. This work was supported by the Max Planck Society and has received funding from the European Research Council under the European Union’s Horizon 2020 research and innovation program (Grant Agreement No. 804824 to J.G.) and from the Deutsche Forschungsgemeinschaft in the Collaborative Research Centre 1080 (to J.G.). Y.K. W. is supported by the Add-on Fellowship of the Joachim Herz Foundation.

## DATA AVAILABILTY

The code used for model simulations is available at GitHub https://github.com/comp-neural-circuits/inhibition-stabilization-paradoxical-effect-STP. All simulation parameters are listed in Supplemental Tables.

## Supplemental Information for

## Supplemental Tables

**Table 1:** IS index.

| Table 1: IS index |  |  |
| --- | --- | --- |
| E-to-E | STD | $\text{Re} \left[ \frac{1}{2} (\tau_E^{-1} (x J_{EE} \alpha_E r_E^{\frac{\alpha_E-1}{\alpha_E}} - 1) - \tau_x^{-1} - U_d r_E) + \sqrt{\frac{1}{4} (\tau_E^{-1} (x J_{EE} \alpha_E r_E^{\frac{\alpha_E-1}{\alpha_E}} - 1) + \tau_x^{-1} + U_d r_E)^2 - \tau_E^{-1} J_{EE} \alpha_E r_E^{\frac{2\alpha_E-1}{\alpha_E}} U_d x} \right]$ |
| | STF | $\text{Re} \left[ \frac{1}{2} (\tau_E^{-1} (u J_{EE} \alpha_E r_E^{\frac{\alpha_E-1}{\alpha_E}} - 1) - \tau_u^{-1} - U_f r_E) + \sqrt{\frac{1}{4} (\tau_E^{-1} (u J_{EE} \alpha_E r_E^{\frac{\alpha_E-1}{\alpha_E}} - 1) + \tau_u^{-1} + U_f r_E)^2 + \tau_E^{-1} J_{EE} \alpha_E r_E^{\frac{2\alpha_E-1}{\alpha_E}} U_f (U_{max} - u)} \right]$ |
| E-to-I | STD | $\tau_E^{-1} (J_{EE} \alpha_E r_E^{\frac{\alpha_E-1}{\alpha_E}} - 1)$ |
| | STF | $\tau_E^{-1} (J_{EE} \alpha_E r_E^{\frac{\alpha_E-1}{\alpha_E}} - 1)$ |
| I-to-E | STD | $\tau_E^{-1} (J_{EE} \alpha_E r_E^{\frac{\alpha_E-1}{\alpha_E}} - 1)$ |
| | STF | $\tau_E^{-1} (J_{EE} \alpha_E r_E^{\frac{\alpha_E-1}{\alpha_E}} - 1)$ |
| I-to-I | STD | $\tau_E^{-1} (J_{EE} \alpha_E r_E^{\frac{\alpha_E-1}{\alpha_E}} - 1)$ |
| | STF | $\tau_E^{-1} (J_{EE} \alpha_E r_E^{\frac{\alpha_E-1}{\alpha_E}} - 1)$ |

**Table 2:** Parameters for Figure 1.

| Table 2: Parameters for Figure 1 |  |  |  |
| --- | --- | --- | --- |
| Symbol | Value | Unit | Description |
| $\tau_E$ | 20 | ms | Time constant of excitatory firing dynamics |
| $\tau_x$ | 200 | ms | Time constant of STD |
| $U_d$ | 1.0 | - | Depression factor |
| Note that these values are also used elsewhere unless mentioned otherwise. |  |  |  |

**Table 3:** Parameters for Figure 2.

| Table 3: Parameters for Figure 2 |  |  |  |
| --- | --- | --- | --- |
| Symbol | Value | Unit | Description |
| $\tau_u$ | 200 | ms | Time constant of STF |
| $U_f$ | 1.0 | - | Facilitation factor |
| $U_{max}$ | 6.0 | - | Maximal facilitation value |
| Note that these values are also used elsewhere unless mentioned otherwise. |  |  |  |

## Supplementary Figures

**Figure S1.**
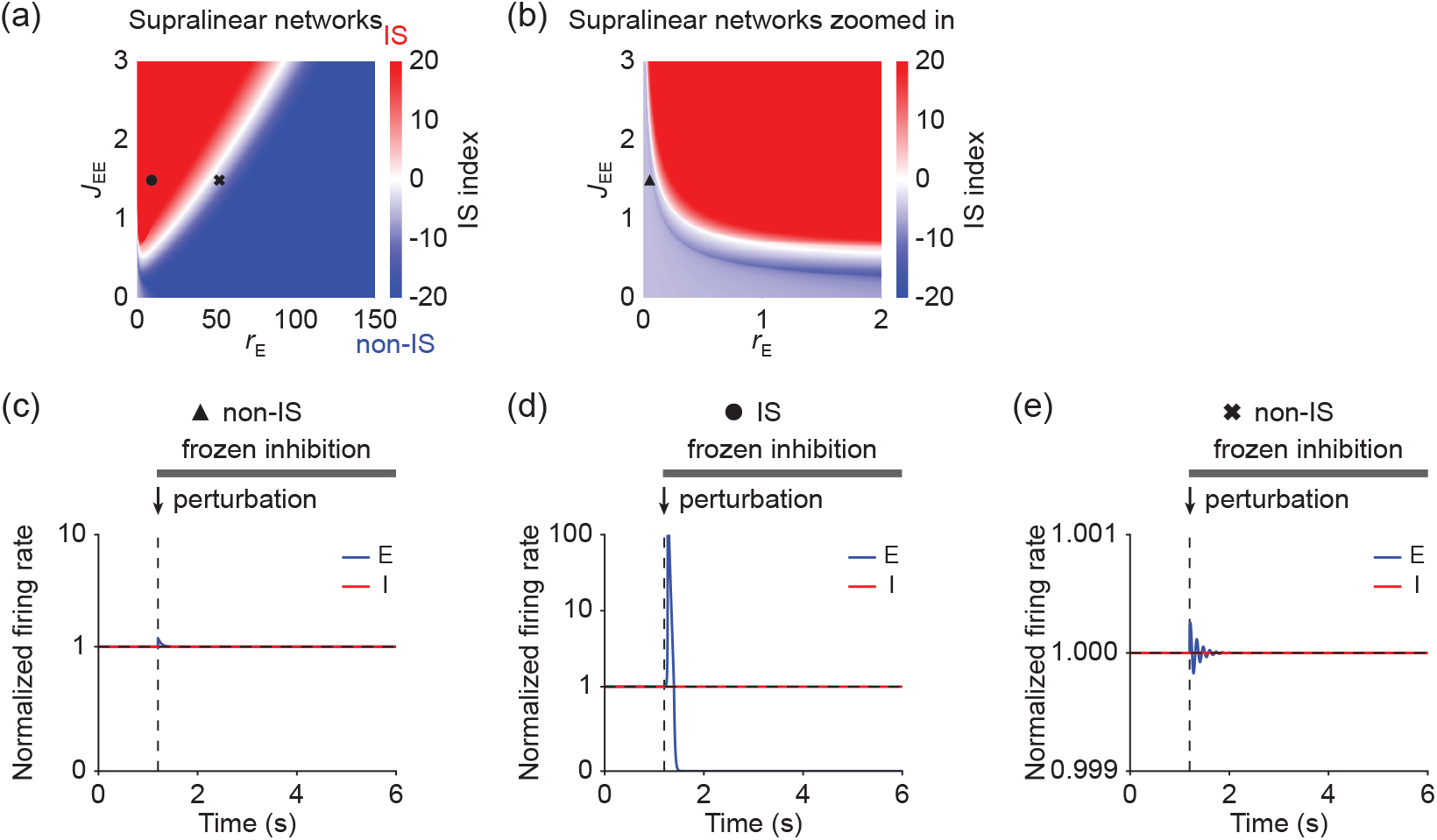
Examples of network activity in response to transient small perturbations to the excitatory activity while freezing inhibition at different initial states. (a) IS index as a function of excitatory firing rate *r*_*E*_ and excitatory-to-excitatory connection strength *J*_*EE*_ for supralinear networks with E-to-E STD. The non-IS and IS regimes are marked in blue and red, respectively. (b) Zoomed-in version of (a). (c) Firing rates of the excitatory (blue) and inhibitory population (red) for a network with an initially non-IS state corresponding to the triangle in (b). A small transient perturbation to excitatory population activity is introduced marked with arrows while freezing inhibition. The periods in which inhibition is frozen are marked with the black bar. (d-e) Same as (c) but for networks with initially IS and non-IS states corresponding to the circle and the cross in (a), respectively. Initial states are shifted by changing *g*_*E*_. *g*_*E*_ is 6.65 in (c), 20 in (d), and 100 in (e).

**Figure S2.**
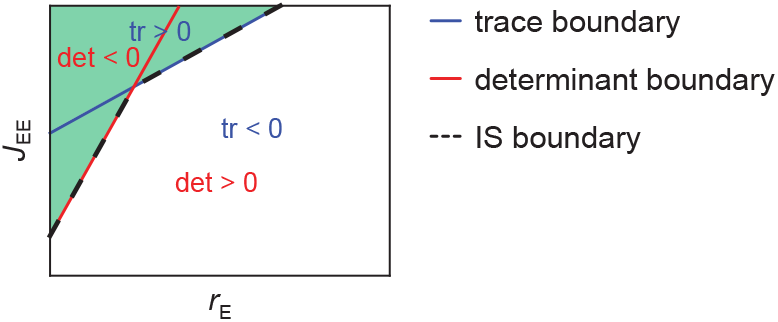
Schematic of IS boundary. Trace boundary (blue line) and determinant boundary (red line) correspond to the *J*_*EE*_ values at which the trace and the determinant of the Jacobian of the system with fixed inhibition are 0 for different *r*_*E*_, respectively. IS boundary (black dashed line) is defined as the minimal *J*_*EE*_ values at which either the trace or the determinant of the Jacobian of the system with fixed inhibition is 0 for different *r*_*E*_. The green and white regions represent the IS regime and non-IS regime, respectively.

**Figure S3.**
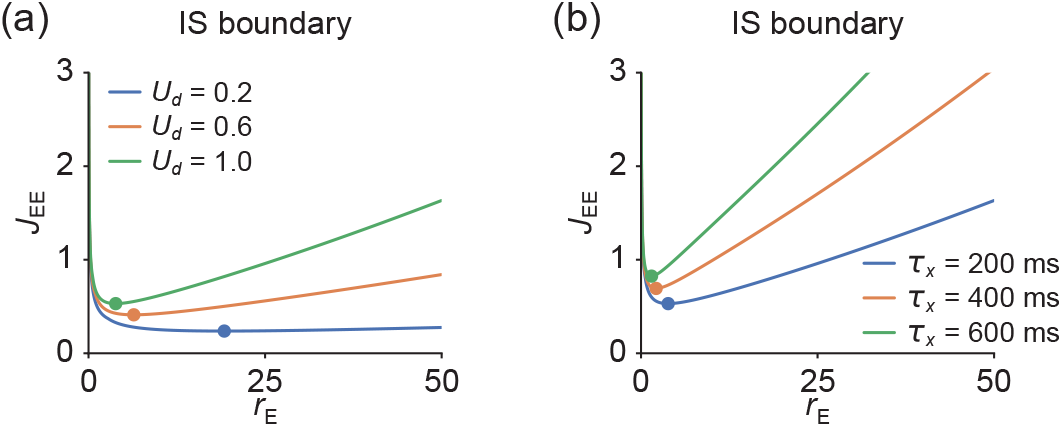
Turning points of the IS boundary in supralinear networks with E-to-E STD. (a) Turning points (dots) of the IS boundary for different values of depression factor *U*_*d*_ represented by different colors. Here, *τ*_*x*_ is 200 ms. (b) Same as (a) but for different values of STD time constant *τ*_*x*_. Here, *U*_*d*_ is 1.

**Figure S4.**
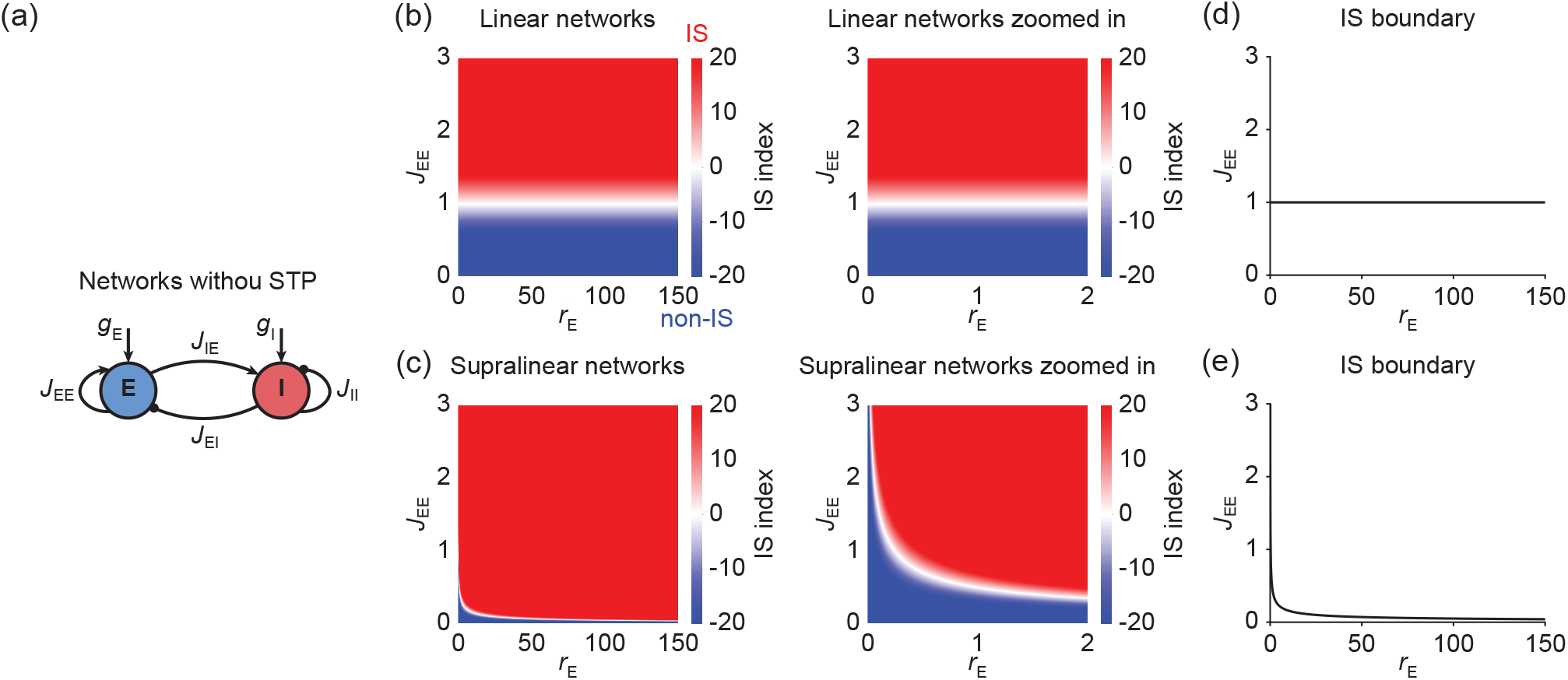
Inhibition stabilization in recurrent neural networks with static connectivity. (a) Schematic of the recurrent network model consisting of an excitatory (blue) and an inhibitory population (red) with static connectivity. (b) Left: IS index as a function of excitatory firing rate *r*_*E*_ and excitatory-to-excitatory connection strength *J*_*EE*_ for linear networks with static connectivity. The non-IS and IS regime are marked in blue and red, respectively. Right: zoomed-in version of (b) left. (c) Same as (b) but for supralinear networks with static connectivity. (d) IS boundary for linear networks with static connectivity, defined as the corresponding excitatory-to-excitatory connection strength 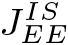 for different *r*_*E*_ at which the IS index is 0. (e) Same as (d) but for supralinear networks with static connectivity.

**Figure S5.**
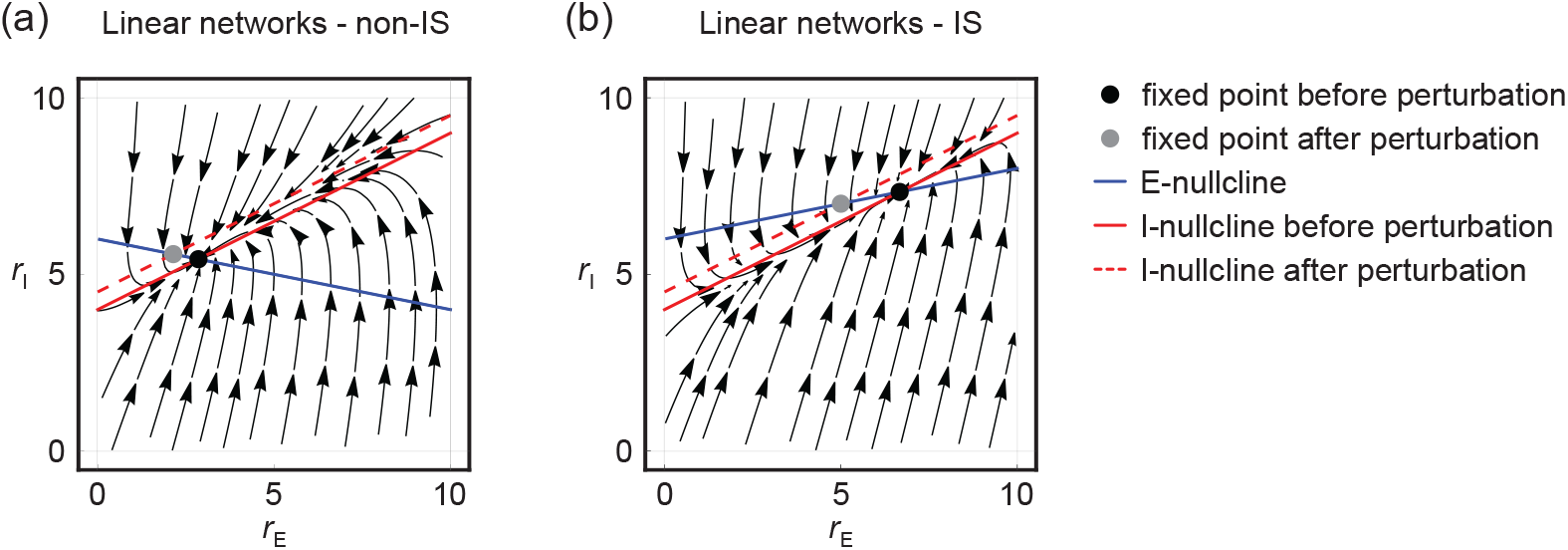
Conditions for the presence of the paradoxical effect from the angle of phase portraits. (a) Phase portrait of a linear non-IS network. The slope of the E-nullcline (blue) around the fixed point (black dot) is negative. The slope of the I-nucllcline (red) is larger than that of the E-nullcline. Perturbating the inhibitory population via injecting additional excitatory current shifts the I-nullcline vertically (from the red solid line to the red dashed line). The inhibitory activity at the new fixed point (gray dot) after perturbation is higher than at the original fixed point. The non-IS network therefore does not exhibit the paradoxical response. (b) Same as (a) but for a linear IS network. The slope of the E-nullcline (blue) around the fixed point is positive. The inhibitory activity at the new fixed point (gray dot) after perturbation is lower than at the original fixed point. The IS network therefore exhibits the paradoxical response.

**Figure S6.**
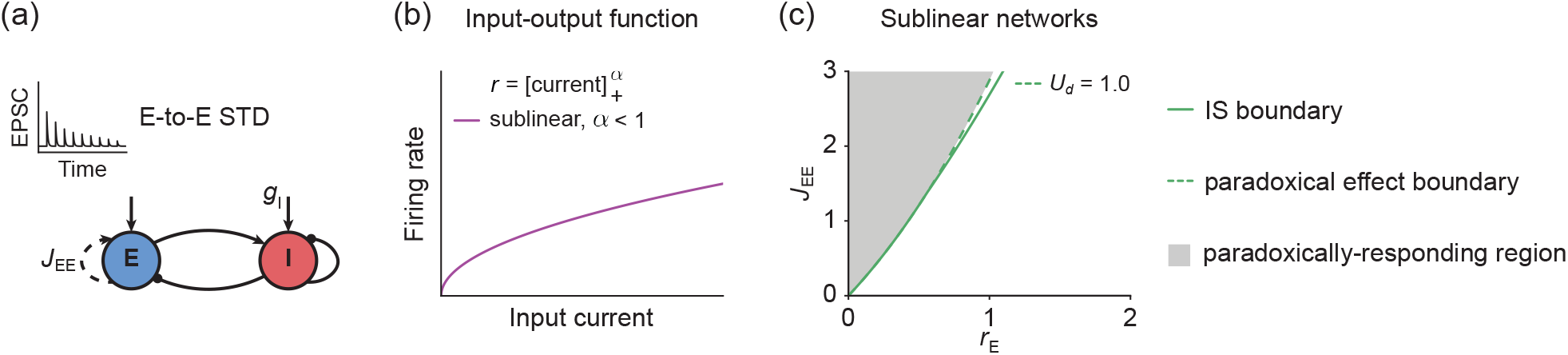
Relationship between inhibition stabilization and the paradoxical effect in sublinear networks with E-to-E STD. (a) Schematic of the recurrent network model consisting of an excitatory (blue) and an inhibitory population (red) with E-to-E STD. (b) Sublinear (purple) input-output function given by a rectified power law with exponent *α* = 0.5. (c) An example of the paradoxical effect boundary (dashed line) and the inhibition stabilization boundary (solid line) as a function of excitatory firing rate *r*_*E*_ for sublinear networks with E-to-E STD. The paradoxically-responding region is marked in gray. Here, *τ*_*x*_ is 200 ms.

**Figure S7.**
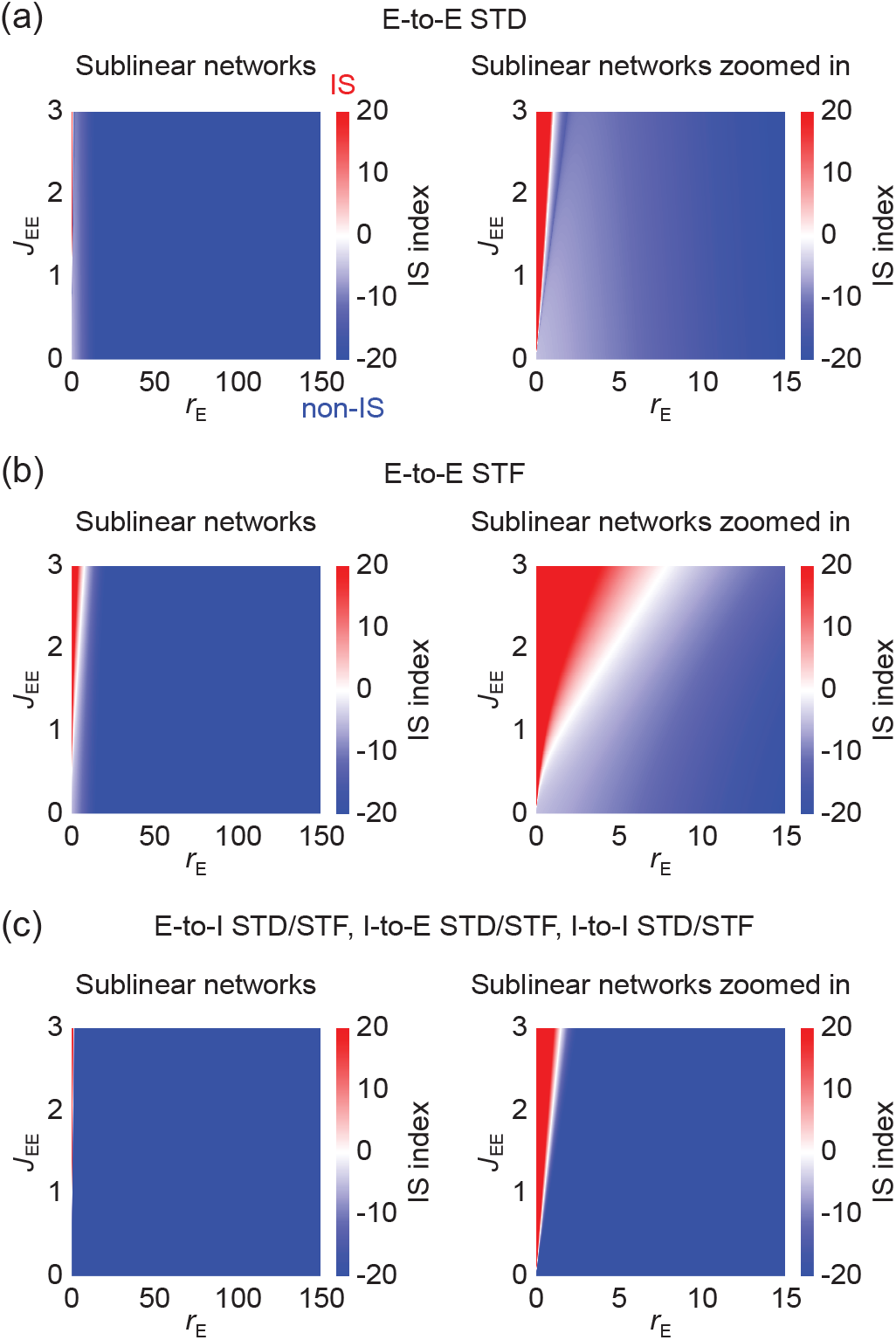
Inhibition stabilization in sublinear networks with short-term plasticity. (a) Left: IS index as a function of excitatory firing rate *r*_*E*_ and excitatory-to-excitatory connection strength *J*_*EE*_ for sublinear networks with E-to-E STD. The non-IS and IS regime are marked in blue and red, respectively. Right: zoomed-in version of (a) left. Here, the depression factor *U*_*d*_ is 1.0, *τ*_*x*_ is 200 ms, the firing rate *r*_*E*_ is plotted in a biologically realistic range from 0 to 150 Hz. (b) Same as (a) but for sublinear networks with E-to-E STF. Here, the facilitation factor *U*_*f*_ is 1.0, *U*_*max*_ is 6.0, and *τ*_*u*_ is 200 ms. (c) Same as (a) but for sublinear networks with E-to-I STD/STF, I-to-E STD/STF, or I-to-I STD/STF.

## Supplemental Methods

To incorporate short-term plasticity mechanisms, we followed the Tsodyks and Markram model [15] and modeled STD as follows:

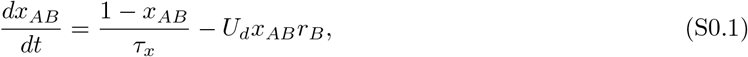

where *x*_*AB*_ is a depression variable that is limited to the interval (0, 1] for the synaptic connection from population *B* to population *A, τ*_*x*_ is the time constant of STD, and *U*_*d*_ is the depression factor.

Furthermore, we modeled STF as follows:

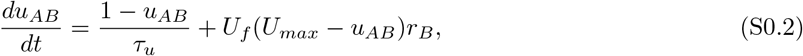

where *u*_*AB*_ is a facilitation variable that is constrained to the interval [1, *U*_*max*_) for the synaptic connection from population *B* to population *A, τ*_*u*_ is the time constant of STF, *U*_*f*_ is the facilitation factor, and *U*_*max*_ is the maximal facilitation value. Note that the short-term plasticity (STP) variables *x*_*AB*_ and *u*_*AB*_ multiply the corresponding synaptic connection strength *J*_*AB*_, and therefore affect the effective synaptic strength.

### Conditions for IS in networks with E-to-E STD

The dynamics of networks with E-to-E STD are given by

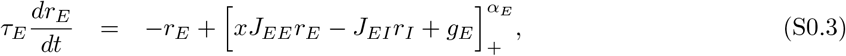

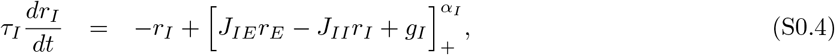

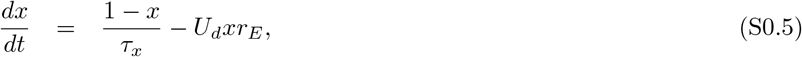

where *x* is the depression variable, which is limited to the interval (0, 1], *τ*_*x*_ is the depression time constant, and *U*_*d*_ is the depression factor.

The Jacobian 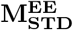 of the system with E-to-E STD is given by

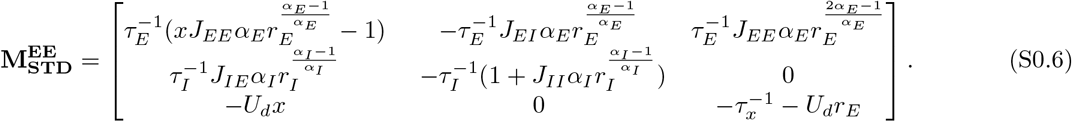

If inhibition is frozen, in other words, if feedback inhibition is absent, the Jacobian of the system becomes as follows:

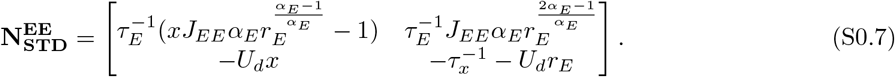

For the system with frozen inhibition, the dynamics are stable if

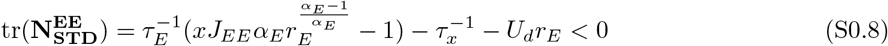

and

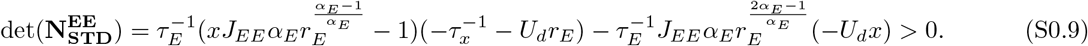

Therefore, if the network is an IS at the fixed point, the following condition has to be satisfied:

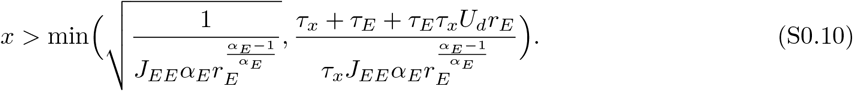

We further define the IS index for the system with E-to-E STD, which is the real part of the leading eigenvalue of 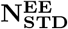, as follows:

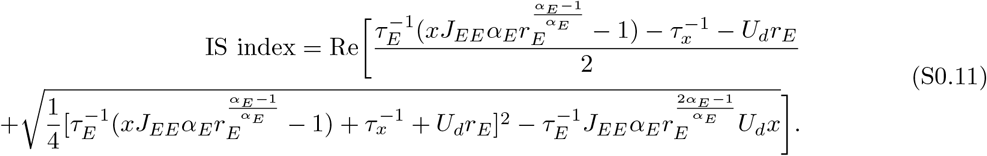

### IS boundary for networks with E-to-E STD

We next investigated how the boundary between non-IS and IS, which we called ‘IS boundary’, changes as a function of *r*_*E*_. Mathematically, the IS boundary is determined by the corresponding recurrent excitatory-to-excitatory connection strength denoted by 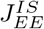 for different *r*_*E*_ at which the IS index is 0. Therefore, we have

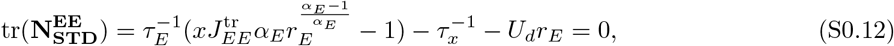

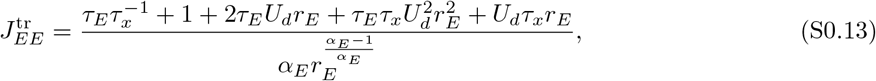

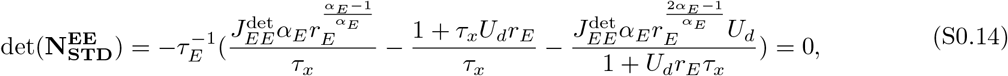

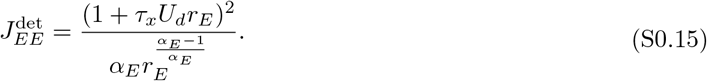

The IS boundary is the minimum of 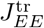 and 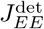 (Fig. S2), and thus can be written as follows:

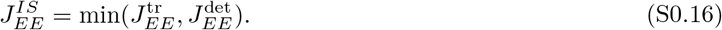

For *α*_*E*_ = 1, we have

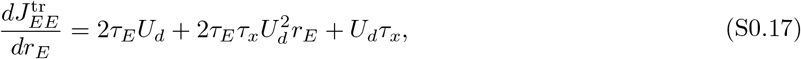

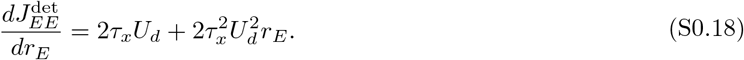

As a result, for linear networks with E-to-E STD, 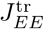 and 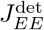 are increasing with *r*_*E*_. Therefore, 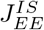 also increases with *r*_*E*_. Furthermore, the increase of 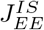 accelerates for larger *r*_*E*_.

For *α*_*E*_ *>* 1, we have

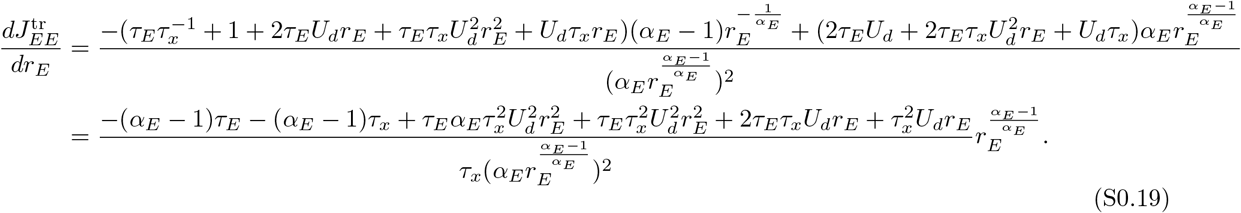

In the small *r*_*E*_ limit, we have:

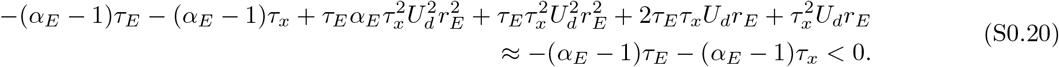

In the large *r*_*E*_ limit, the increasing neural gain-related term begins to dominate, and we have:

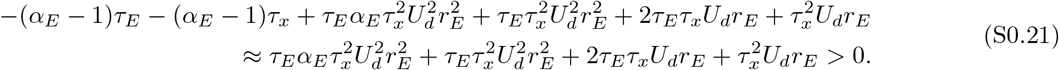

Therefore, 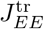 first decreases and then increases with *r*_*E*_.

Similarly, we have:

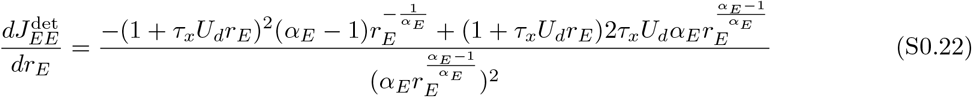

and

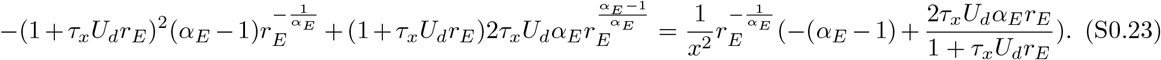

In the small *r*_*E*_ limit, the short-term depression variable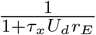 is close to 1, whereas the neural gain-related term 2*τ*_*x*_*U*_*d*_*α*_*E*_*r*_*E*_ is close to 0, so we have:

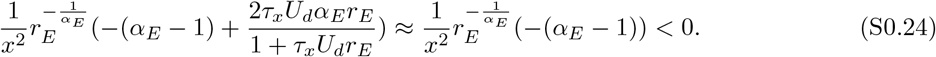

As *r*_*E*_ increases, the short-term depression variable 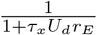 tends to decrease the expression 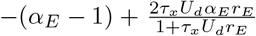, whereas the increasing neural gain-related term 2*τ*_*x*_*U*_*d*_*α*_*E*_*r*_*E*_ tends to increase the expression 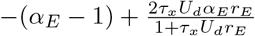. In the large *r*_*E*_ limit, we have:

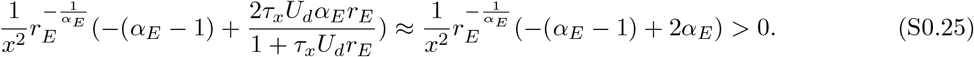

Therefore, 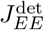 first decreases and then increases with *r*_*E*_.

In sum, for supralinear networks with E-to-E STD, the IS boundary changes non-monotonically with *r*_*E*_. More specifically, it first shifts downwards and then shifts upwards as *r*_*E*_ increases.

For *α*_*E*_ *<* 1, we have

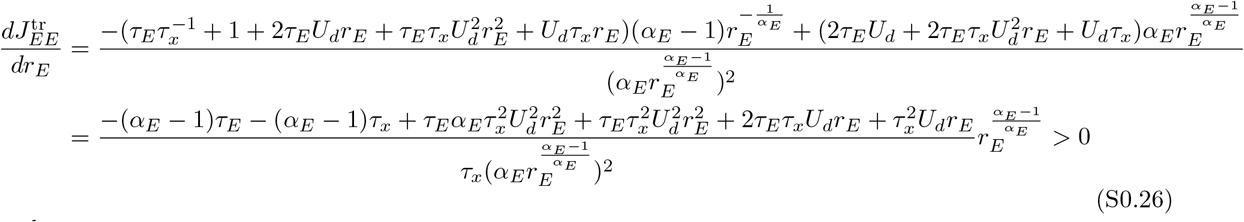

and

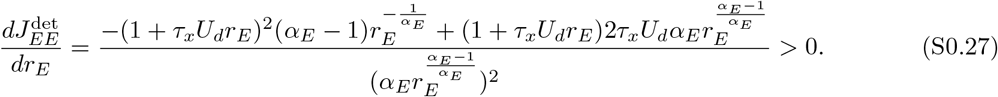

Therefore, for sublinear networks with E-to-E STD, the IS boundary changes monotonically with *r*_*E*_. More specifically, it always shifts upwards as *r*_*E*_ increases.

### Turning point of the IS boundary for networks with E-to-E

Here, we sought to investigate how the turning point of the IS boundary changes with the short-term depression factor *U*_*d*_ and the short-term depression time constant *τ*_*x*_. Assuming that around the turning point, 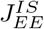 is some smooth function of *r*_*E*_, the turning point 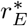 of the IS boundary can then be defined as follows:

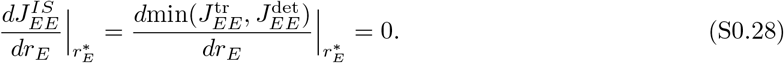

Furthermore, the turning point 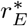 of the IS boundary can be described as the minimum of the turning point 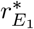of 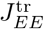 and the turning point 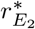 of 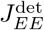, i.e.,

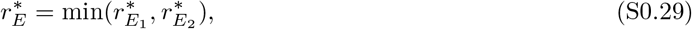

with

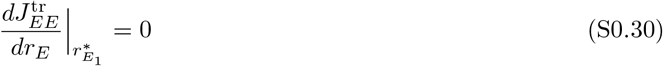

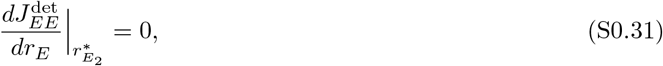

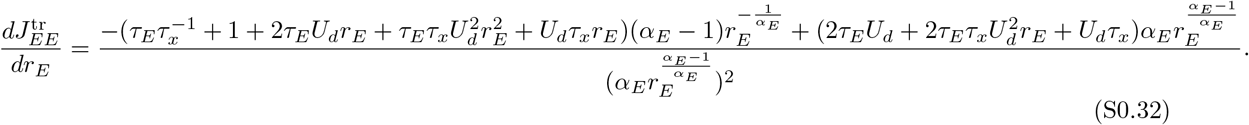

Therefore, we have

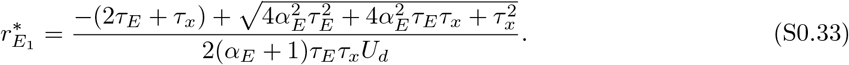

The turning point 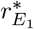 of 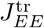 can therefore be expressed as a function of *U*_*d*_ and *τ*_*x*_. To identify how *U*_*d*_ and *τ*_*x*_ affect 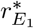, we computed the following derivatives:

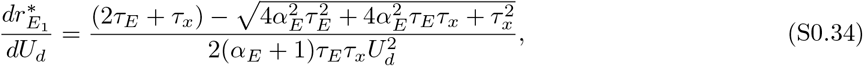

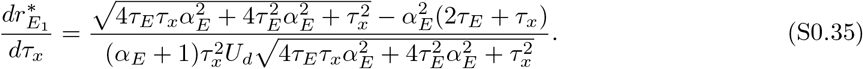

For *α >* 1, we have:

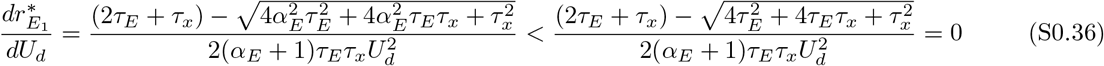

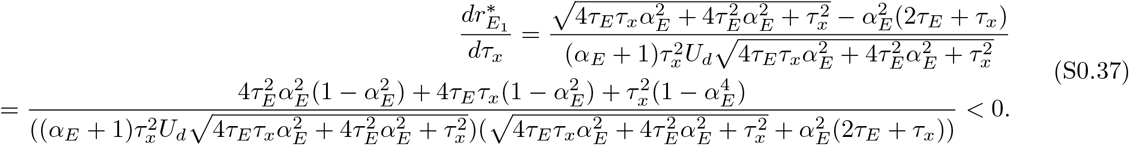

Therefore, for *α >* 1, increasing either *U*_*d*_ or *τ*_*x*_ shifts the turning point 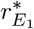 of 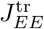 to the left in the coordinate system in which the abscissa is *r*_*E*_ and the ordinate is *J*_*EE*_. Furthermore,

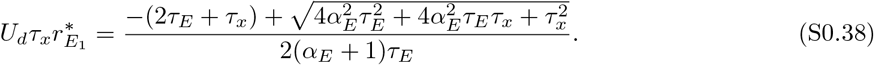

When *α*_*E*_ is of order 1, then 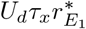 is approximately of the same order as 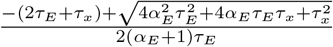 which is of order 1. Therefore, the turning point 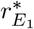 of 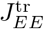 appears when 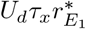 is of order 1.

Similarly, we have:

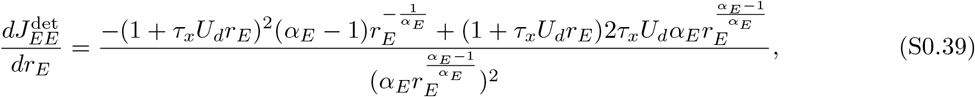

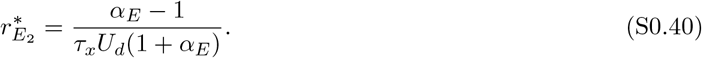

Therefore, for *α >* 1, increasing either *U*_*d*_ or *τ*_*x*_ shifts the turning point 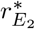 of 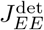 to the left in the coordinate system in which the abscissa is *r*_*E*_ and the ordinate is *J*_*EE*_. Moreover,

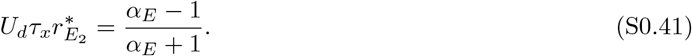

Therefore, the turning point 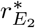 of 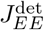 appears when 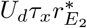 is of order 1.

As the turning point 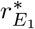 of 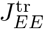 and the turning point 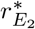 of 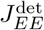 are affected by *U*_*d*_ and *τ*_*x*_ in the same way, increasing either *U*_*d*_ or *τ*_*x*_ also shifts the turning point 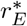 of the IS boundary to the left in the same coordinate system. Furthermore, both 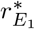 and 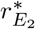 appear when 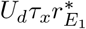 and 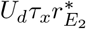 are of order 1, respectively, the turning point 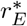 of the IS boundary therefore appears when 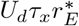 is of order 1.

In addition, since

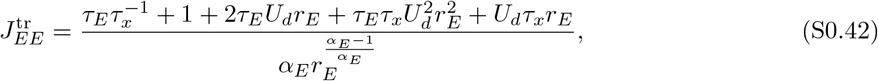

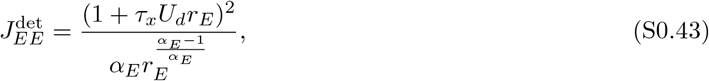

it is obvious that increasing *U*_*d*_ shifts the turning point 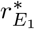up in the (*r*_*E*_, *J*_*EE*_) coordinate system, and increasing *U*_*d*_ or *τ*_*x*_ shifts the turning point 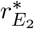 up. For fixed *r*_*E*_, we have:

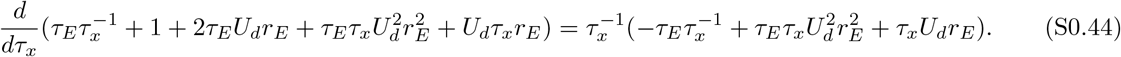

The rate dynamics is typically an order of magnitude larger than the dynamics of the short-term plasticity mechanisms. As we derived before, at the turning point, *τ*_*x*_*U*_*d*_*r*_*E*_ is of order 1. So the above expression is positive in biologically realistic regimes, indicating that increasing *τ*_*x*_ shifts the turning point 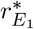 up in the (*r*_*E*_, *J*_*EE*_) coordinate system. Therefore, increasing either *U*_*d*_ or *τ*_*x*_ also shifts the turning point 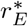 of the IS boundary up in that coordinate system.

### Conditions for paradoxical response in networks with E-to-E STD

Next, we derived the condition of having the paradoxical effect in networks with E-to-E STD. Under the assumption that sufficient small perturbations would not lead to unstable network dynamics and regime transition between non-IS and IS, the conditions of having paradoxical effects are a positive slope of the excitatory nullcline and a larger slope of the inhibitory nullcline than that of the excitatory nullcline locally around the fixed point [17]. We can write the excitatory nullcline as follows

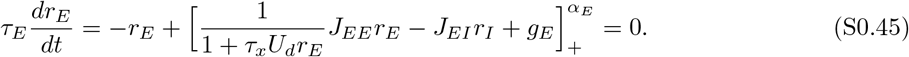

For *r*_*E*,*I*_ *>* 0, we have

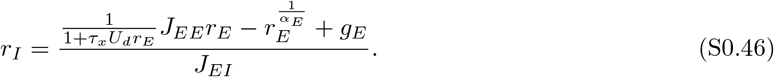

The slope of the excitatory nullcline in the *r*_*E*_*/r*_*I*_ plane where *x* axis is *r*_*E*_ and *y* axis is *r*_*I*_ can be written as follows

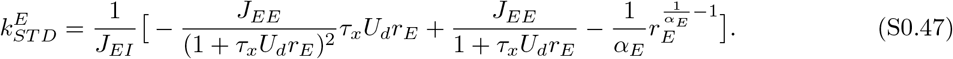

Note that as the short-term depression variable *x*_*EE*_ depends on *r*_*E*_ and changes dynamically, the slope of the excitatory nullcline 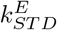 also contains the term 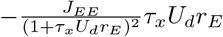, which is the product of *J*_*EE*_*r*_*E*_ and the derivative of short-term depression variable *x*_*EE*_ with respect to *r*_*E*_.

To have paradoxical effect, the slope of the excitatory nullcline at the fixed point of the system has to be positive. Therefore, the short-term depression variable *x* at the fixed point has to satisfy the following condition

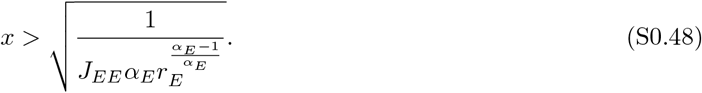

The inhibitory nullcline can be written as follows

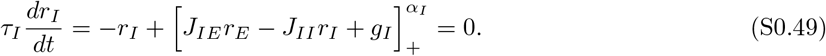

In the region of rates *r*_*E*,*I*_ *>* 0, we have

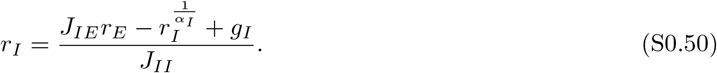

The slope of the inhibitory nullcline can be written as follows

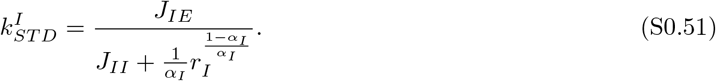

In addition to the positive slope of the excitatory nullcline, the slope of the inhibitory nullcline at the fixed point of the system has to be larger than the slope of the excitatory nullcline. We therefore have

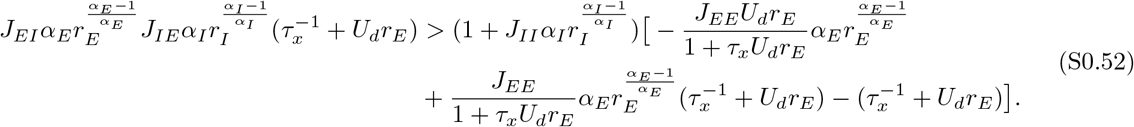

Note that to ensure that the system with E-to-E STD is stable, 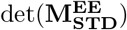 has to be negative. Therefore, we have

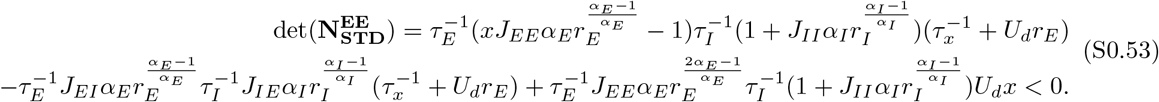

The condition shown in Eq. S0.52 is the same as the stability condition of the determinant of the Jacobian of the system with E-to-E STD, namely, as 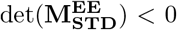. Thus, the condition is always satisfied when the system with E-to-E STD is stable.

Based on the condition of being IS shown in Eq. S0.10 and the condition of having paradoxical effect shown in Eq. S0.48, we therefore can conclude that in networks with E-to-E STD, the paradoxical effect implies inhibitory stabilization, whereas inhibitory stabilization does not necessarily imply paradoxical responses.

Furthermore, for networks with E-to-E STD exhibiting no paradoxical effects, 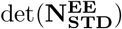 is larger than 0, indicating that the excitatory nullcline has a negative slope around the fixed point in the *r*_*E*_/*r*_*I*_ plane where *x* axis is *r*_*E*_ and *y* axis is *r*_*I*_ . However, the networks can be inhibition stabilized when

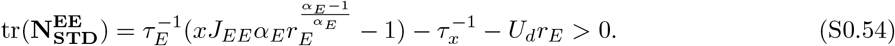

Then, 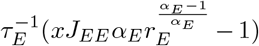 is also positive, suggesting that the excitatory subnetwork is unstable if STD variables and inhibition are fixed. Therefore, in the presence of E-to-E STD, networks in which dynamic STD regulation is required to ensure stability as studied in [18, 19] are a subset of inhibition stabilized networks which do not exhibit paradoxical effects.

### Paradoxical boundary for networks with E-to-E STD

We next investigated how the boundary between non-paradoxical and paradoxical effects, which we called ‘paradoxical boundary’, changes as a function of *r*_*E*_. Mathematically, the paradoxical boundary is determined by the corresponding recurrent excitatory-to-excitatory connection strength denoted by 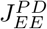 for different *r*_*E*_ at which the slope of the excitatory nullcline is 0. Therefore, we have

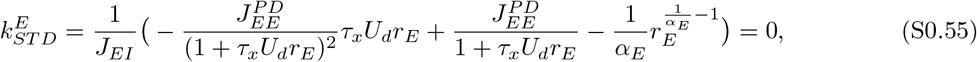

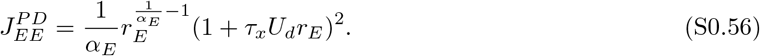

For *α*_*E*_ = 1, we have

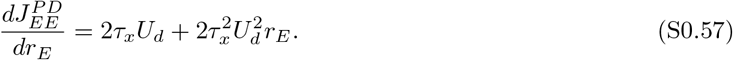

Obviously, the derivative is always positive. Therefore, for linear networks with E-to-E STD, 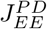 is increasing with *r*_*E*_.

For *α*_*E*_ *>* 1, we have

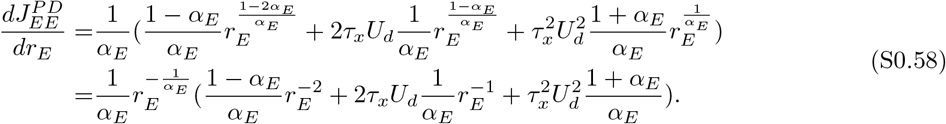

Clearly, the derivative switches from negative to positive as *r*_*E*_ grows. Therefore, for supralinear networks with E-to-E STD, the paradoxical effect boundary first shifts downwards and then shifts upwards as *r*_*E*_ increases.

For *α*_*E*_ *<* 1, we have

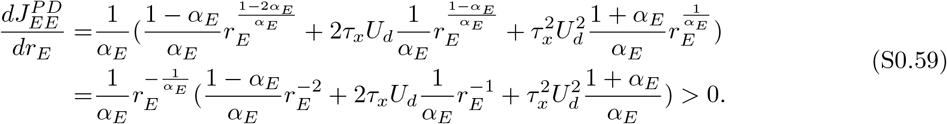

Therefore, for sublinear networks with E-to-E STD, the paradoxical effect boundary always shifts upwards as *r*_*E*_ increases.

### Conditions for IS in networks with E-to-E STF

The dynamics of networks with E-to-E STF are given by

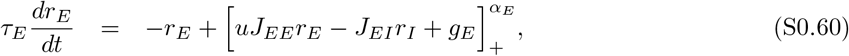

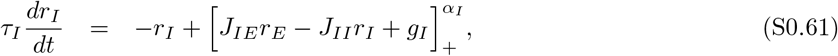

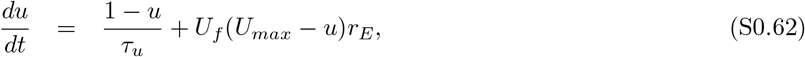

where *u* is the facilitation variable constrained to the interval [1, *U*_*max*_), *U*_*max*_ is the maximal facilitation value, *τ*_*u*_ is the time constant of STF, and *U*_*f*_ is the facilitation factor.

The Jacobian 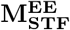 of the system with E-to-E STF is given by

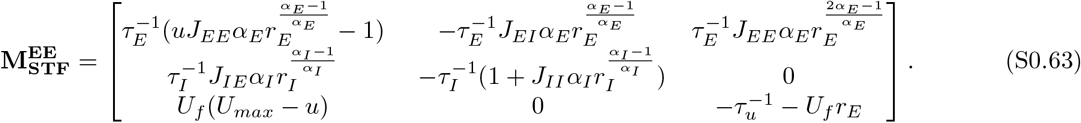

If inhibition is frozen, in other words, if feedback inhibition is absent, the Jacobian of the system becomes as follows:

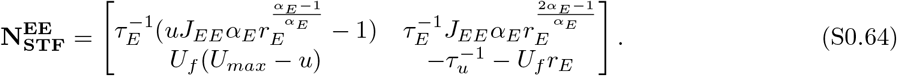

For the system with frozen inhibition, the dynamics are stable if

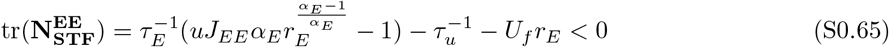

and

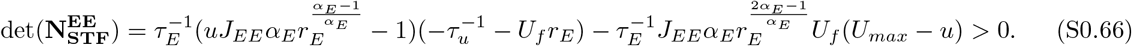

Therefore, if the network is an IS at the fixed point, the following condition has to be satisfied:

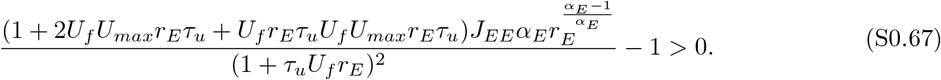

We further define the IS index for the system with E-to-E STF as follows:

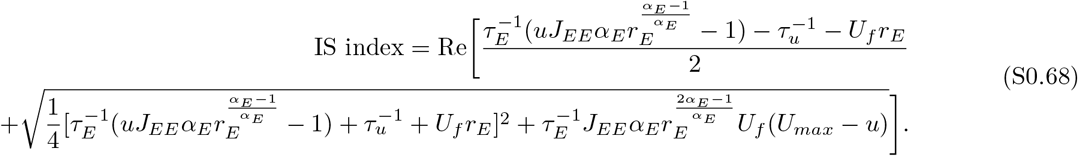

### IS boundary for networks with E-to-E STF

We next investigated how the boundary between non-IS and IS, which we called ‘IS boundary’, changes as a function of *r*_*E*_. Mathematically, the IS boundary is determined by the corresponding recurrent excitatory-to-excitatory connection strength denoted by 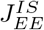 for different *r*_*E*_ at which the IS index is 0. Therefore, we

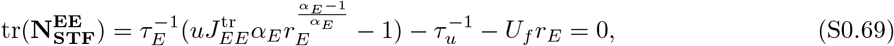

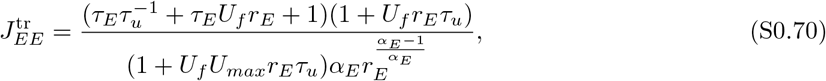

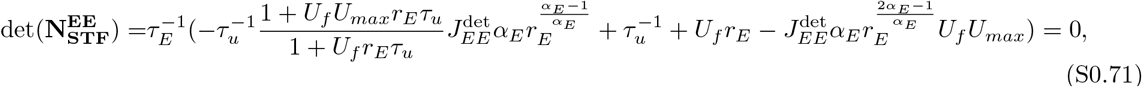

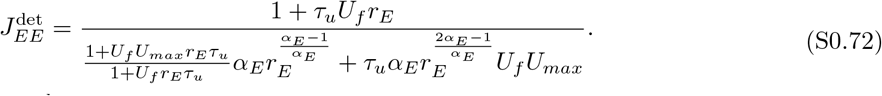

Clearly, 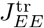 is greater than 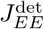 for any *r*_*E*_. Therefore, we have

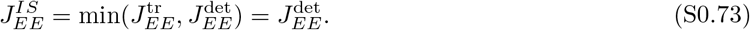

For *α*_*E*_ = 1, we have

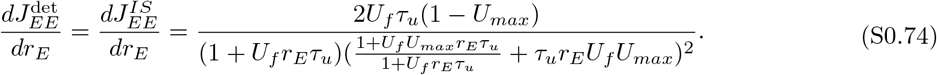

Therefore, for linear networks with E-to-E STF,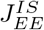 is decreasing with *r*_*E*_. Furthermore, the decrease of 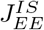 slows down for larger *r*_*E*_.

For *α*_*E*_ *>* 1, we have

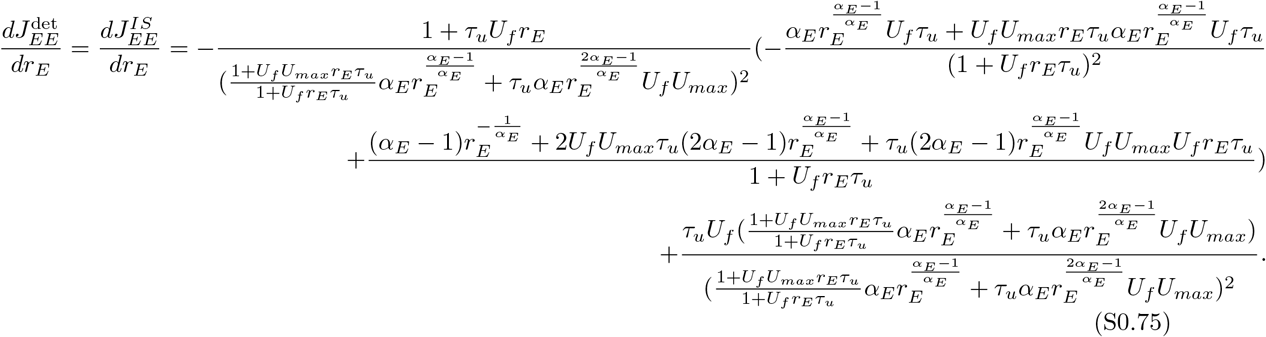

Further,

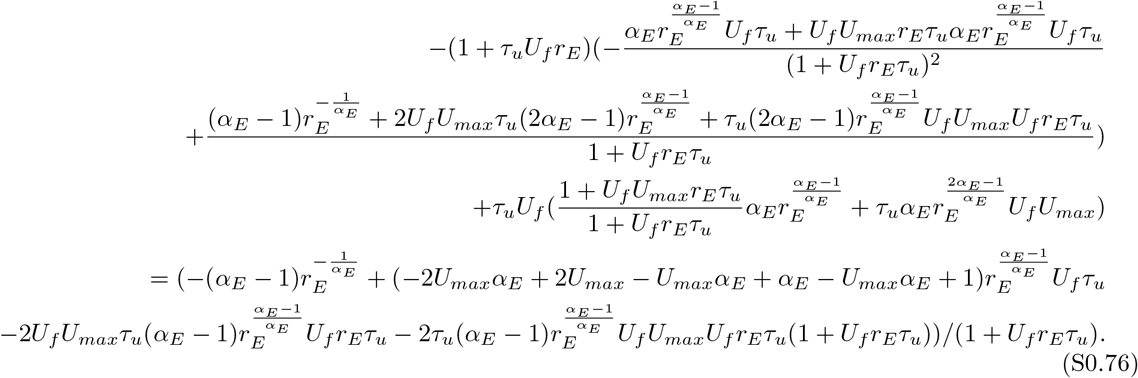

Therefore, for supralinear networks with E-to-E STF, 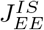 is decreasing with *r*_*E*_.

### Conditions for paradoxical response in networks with E-to-E STF

Next, we derived the condition of having the paradoxical effect in networks with E-to-E STF. Under the assumption that sufficient small perturbations would not lead to unstable network dynamics and regime transition between non-IS and IS, the conditions of having paradoxical effects are a positive slope of the excitatory nullcline and a larger slope of the inhibitory nullcline than that of the excitatory nullcline locally around the fixed point [17]. We can write the excitatory nullcline as follows

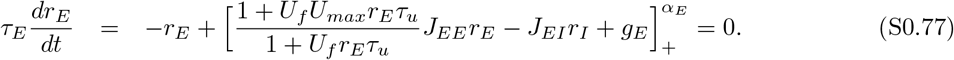

For *r*_*E*,*I*_ *>* 0, we have

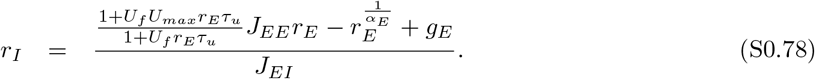

The slope of the excitatory nullcline in the *r*_*E*_*/r*_*I*_ plane where *x* axis is *r*_*E*_ and *y* axis is *r*_*I*_ can be written as follows

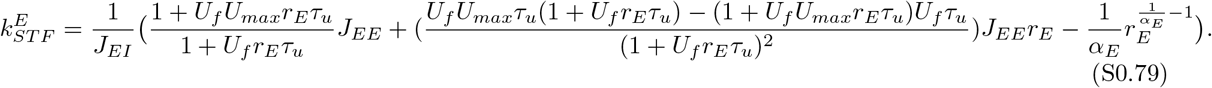

To have paradoxical effect, the slope of the excitatory nullcline at the fixed point of the system has to be positive. Therefore, we have

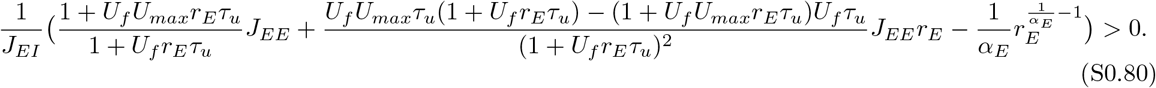

The above condition can be simplified as follows:

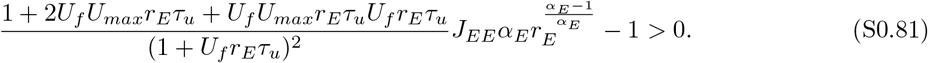

The inhibitory nullcline can be written as follows

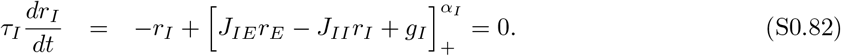

In the region of rates *r*_*E*,*I*_ *>* 0, we have

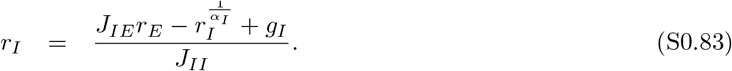

The slope of the inhibitory nullcline can be written as follows

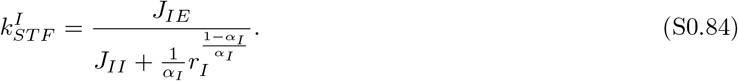

In addition to the positive slope of the excitatory nullcline, the slope of the inhibitory nullcline at the fixed point of the system has to be larger than the slope of the excitatory nullcline. We therefore have

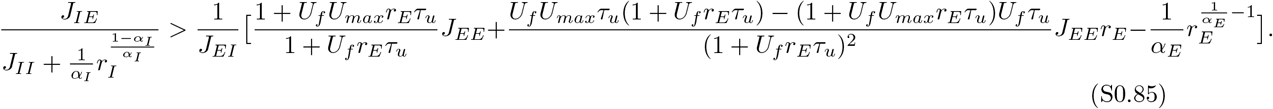

Note that to ensure that the system with E-to-E STF is stable, 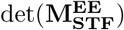 has to be negative. Therefore, we have

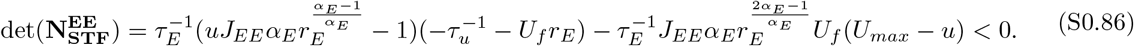

The condition shown in Eq. S0.85 is the same as the stability condition of the determinant of the Jacobian of the system with E-to-E STF, namely, as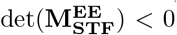 . Thus, the condition is always satisfied when the system with E-to-E STF is stable.

The condition of being IS shown in Eq. S0.67 is identical to the condition of having paradoxical effect shown in Eq. S0.81, we therefore can conclude that in networks with E-to-E STF, inhibitory stabilization and the paradoxical effect imply each other.

### Conditions for IS in networks with E-to-I STD

The dynamics of networks with E-to-I STD are given by

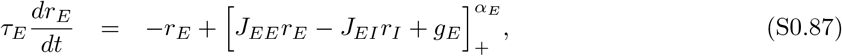

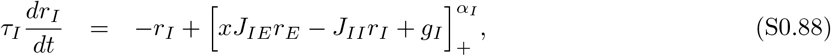

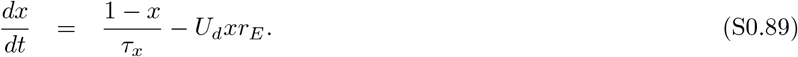

where *x* is the depression variable, which is limited to the interval (0, 1], *τ*_*x*_ is the depression time constant, and *U*_*d*_ is the depression factor.

The Jacobian 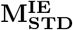 of the system with E-to-I STD is given by

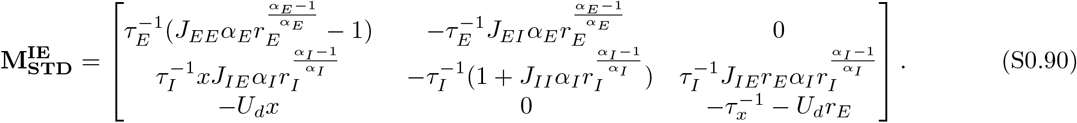

If inhibition is frozen, in other words, if feedback inhibition is absent, the Jacobian of the system becomes as follows:

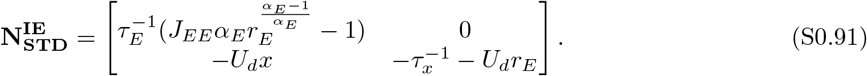

For the system with frozen inhibition, the dynamics are stable if

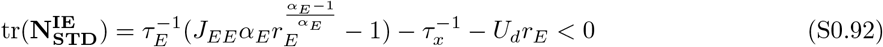

and

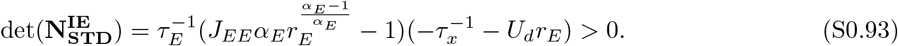

Therefore, if the network is an IS at the fixed point, the following condition has to be satisfied:

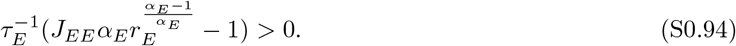

We further define the IS index for the system with E-to-I STD as follows:

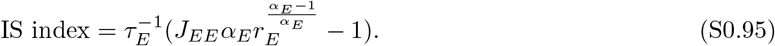

### Conditions for paradoxical response in networks with E-to-I STD

Next, we derived the condition of having the paradoxical effect in networks with E-to-I STD. Under the assumption that sufficient small perturbations would not lead to unstable network dynamics and regime transition between non-IS and IS, the conditions of having paradoxical effects are a positive slope of the excitatory nullcline and a larger slope of the inhibitory nullcline than that of the excitatory nullcline locally around the fixed point [17]. We can write the excitatory nullcline as follows

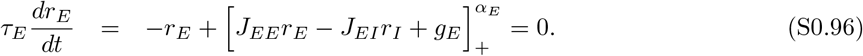

For *r*_*E*,*I*_ *>* 0, we have

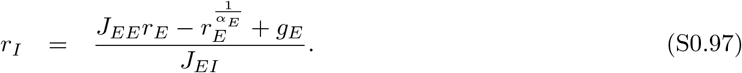

The slope of the excitatory nullcline in the *r*_*E*_*/r*_*I*_ plane where *x* axis is *r*_*E*_ and *y* axis is *r*_*I*_ can be written as follows

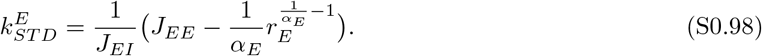

To have paradoxical effect, the slope of the excitatory nullcline at the fixed point of the system has to be positive. Therefore, we have

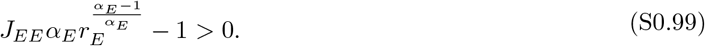

The inhibitory nullcline can be written as follows

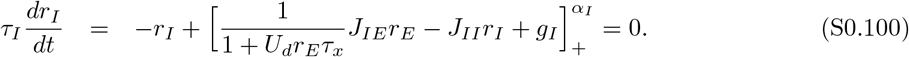

In the region of rates *r*_*E*,*I*_ *>* 0, we have

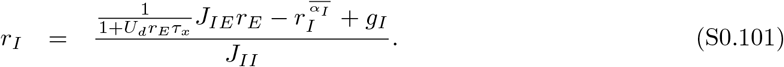

The slope of the inhibitory nullcline can be written as follows

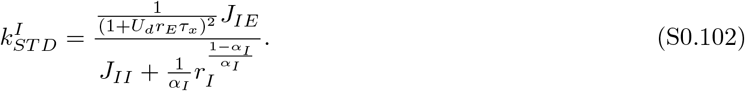

In addition to the positive slope of the excitatory nullcline, the slope of the inhibitory nullcline at the fixed point of the system has to be larger than the slope of the excitatory nullcline. We therefore have

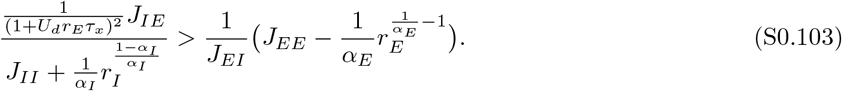

Note that to ensure that the system with E-to-I STD is stable, 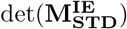 has to be negative. Therefore, we have

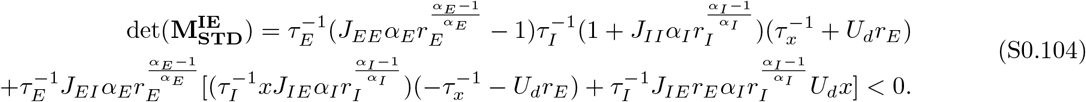

The condition shown in Eq. S0.103 is fulfilled if 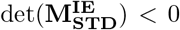. Thus, the condition is always satisfied when the system with E-to-I STD is stable.

The condition of being IS shown in Eq. S0.94 is identical to the condition of having paradoxical effect shown in Eq. S0.99, we therefore can conclude that in networks with E-to-I STD, inhibitory stabilization and the paradoxical effect imply each other.

### Conditions for IS in networks with E-to-I STF

The dynamics of networks with E-to-I STF are given by

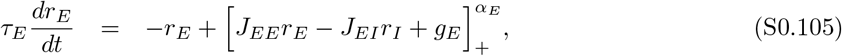

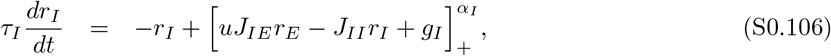

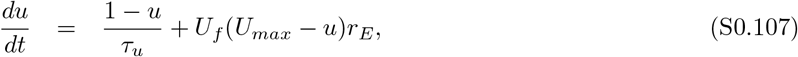

where *u* is the facilitation variable constrained to the interval [1, *U*_*max*_), *U*_*max*_ is the maximal facilitation value, *τ*_*u*_ is the time constant of STF, and *U*_*f*_ is the facilitation factor.

The Jacobian 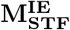 of the system with E-to-I STF is given by

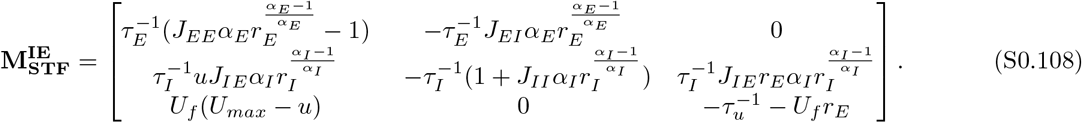

If inhibition is frozen, in other words, if feedback inhibition is absent, the Jacobian of the system becomes as follows:

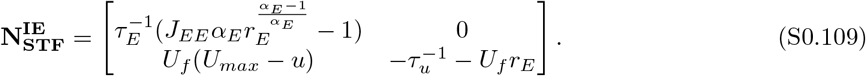

For the system with frozen inhibition, the dynamics are stable if

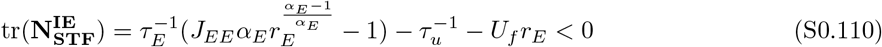

and

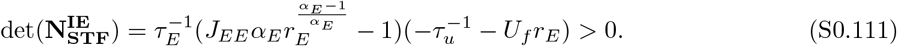

Therefore, if the network is an IS at the fixed point, the following condition has to be satisfied:

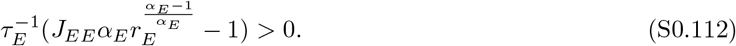

We further define the IS index for the system with E-to-I STF as follows:

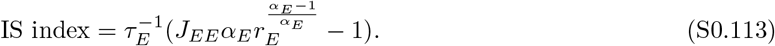

### Conditions for paradoxical response in networks with E-to-I STF

Next, we derived the condition of having the paradoxical effect in networks with E-to-I STF. Under the assumption that sufficient small perturbations would not lead to unstable network dynamics and regime transition between non-IS and IS, the conditions of having paradoxical effects are a positive slope of the excitatory nullcline and a larger slope of the inhibitory nullcline than that of the excitatory nullcline locally around the fixed point [17]. We can write the excitatory nullcline as follows

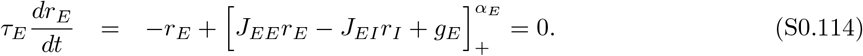

For *r*_*E*,*I*_ *>* 0, we have

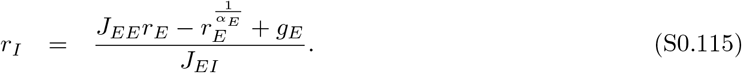

The slope of the excitatory nullcline in the *r*_*E*_*/r*_*I*_ plane where *x* axis is *r*_*E*_ and *y* axis is *r*_*I*_ can be written as follows

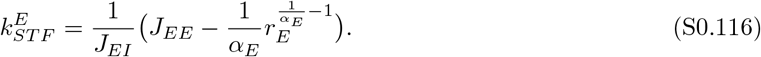

To have paradoxical effect, the slope of the excitatory nullcline at the fixed point of the system has to be positive. Therefore, we have

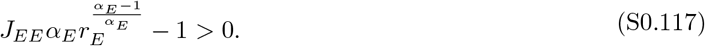

The inhibitory nullcline can be written as follows

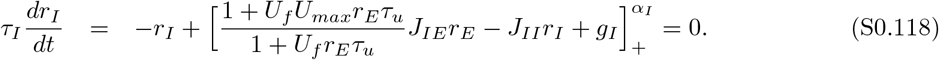

In the region of rates *r*_*E*,*I*_ *>* 0, we have

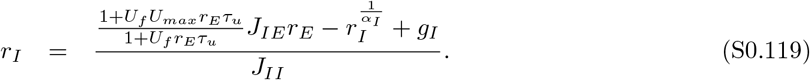

The slope of the inhibitory nullcline can be written as follows

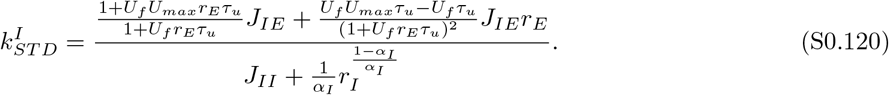

In addition to the positive slope of the excitatory nullcline, the slope of the inhibitory nullcline at the fixed point of the system has to be larger than the slope of the excitatory nullcline. We therefore have

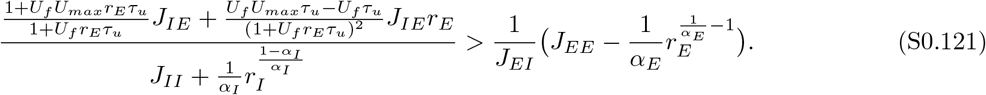

Note that to ensure that the system with E-to-I STF is stable, 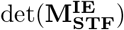 has to be negative. Therefore, we have

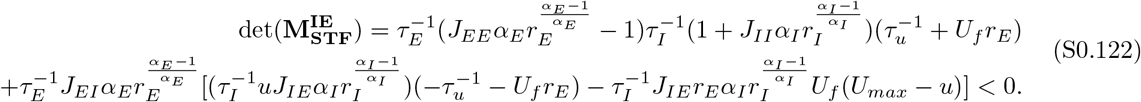

The condition shown in Eq. S0.121 is the same as the stability condition of the determinant of the Jacobian of the system with E-to-I STF, namely, as 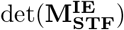. Thus, the condition is always satisfied when the system with E-to-I STF is stable.

The condition of being IS shown in Eq. S0.112 is identical to the condition of having paradoxical effect shown in Eq. S0.117, we therefore can conclude that in networks with E-to-I STF, inhibitory stabilization and the paradoxical effect imply each other.

### Conditions for IS in networks with I-to-E STD

The dynamics of networks with I-to-E STD are given by

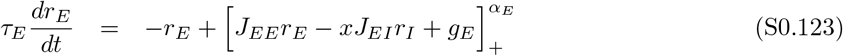

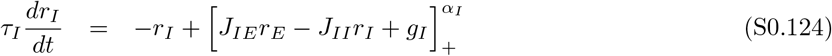

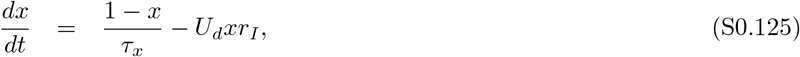

where *x* is the depression variable, which is limited to the interval (0, 1], *τ*_*x*_ is the depression time constant, and *U*_*d*_ is the depression factor.

The Jacobian 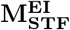 of the system with I-to-E STD is given by

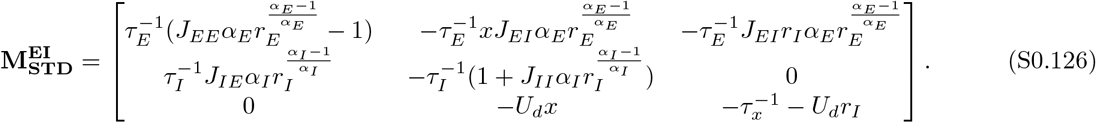

If inhibition is frozen, in other words, if feedback inhibition is absent, the Jacobian of the system becomes as follows:

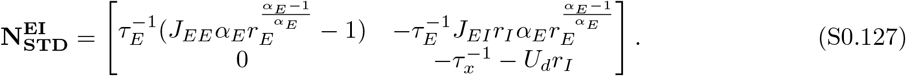

For the system with frozen inhibition, the dynamics are stable if

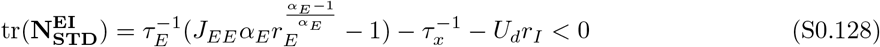

and

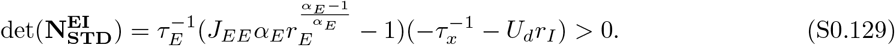

Therefore, if the network is an IS at the fixed point, the following condition has to be satisfied:

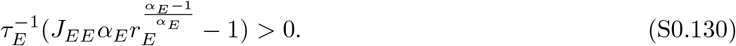

We further define the IS index for the system with I-to-E STD as follows:

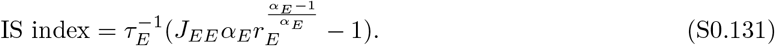

### Conditions for paradoxical response in networks with I-to-E STD

Next, we derived the condition of having the paradoxical effect in networks with I-to-E STD. Under the assumption that sufficient small perturbations would not lead to unstable network dynamics and regime transition between non-IS and IS, the conditions of having paradoxical effects are a positive slope of the excitatory nullcline and a larger slope of the inhibitory nullcline than that of the excitatory nullcline locally around the fixed point [17]. We can write the excitatory nullcline as follows

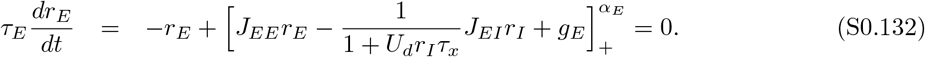

For *r*_*E*,*I*_ *>* 0, we have

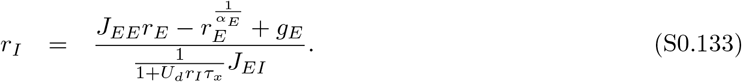

The slope of the excitatory nullcline in the *r*_*E*_*/r*_*I*_ plane where *x* axis is *r*_*E*_ and *y* axis is *r*_*I*_ can be written as follows

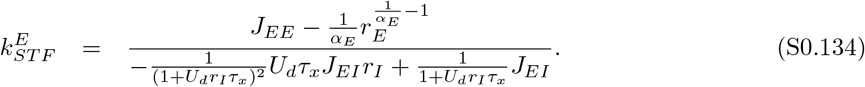

Provided that the inhibitory current to the excitatory network is always an increasing function (at least locally around the fixed point) of the inhibitory firing rate, we have

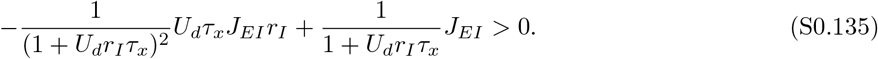

To have paradoxical effect, the slope of the excitatory nullcline at the fixed point of the system has to be positive. Therefore, we have

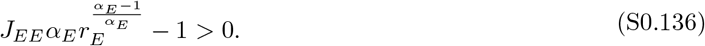

The inhibitory nullcline can be written as follows

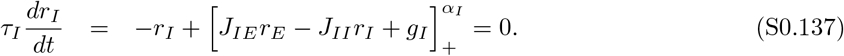

In the region of rates *r*_*E*,*I*_ *>* 0, we have

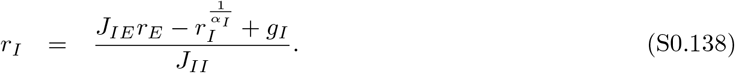

The slope of the inhibitory nullcline can be written as follows

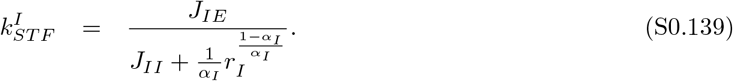

In addition to the positive slope of the excitatory nullcline, the slope of the inhibitory nullcline at the fixed point of the system has to be larger than the slope of the excitatory nullcline. We therefore have

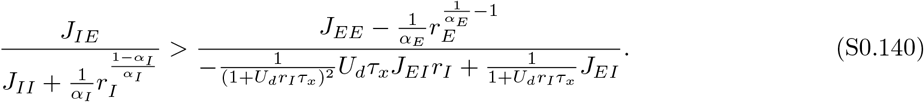

Note that to ensure that the system with I-to-E STD is stable, 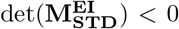 has to be negative. Therefore, we have

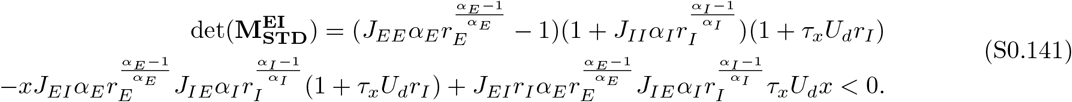

The condition shown in Eq. S0.140 is the same as the stability condition of the determinant of the Jacobian of the system with I-to-E STD, namely, as det 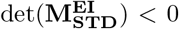. Thus, the condition is always satisfied when the system with I-to-E STD is stable.

The condition of being IS shown in Eq. S0.130 is identical to the condition of having paradoxical effect shown in Eq. S0.136, we therefore can conclude that in networks with I-to-E STD, inhibitory stabilization and the paradoxical effect imply each other.

### Conditions for IS in networks with I-to-E STF

The dynamics of networks with I-to-E STF are given by

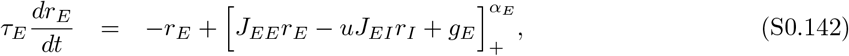

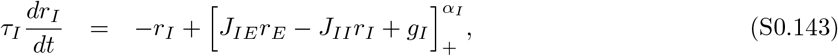

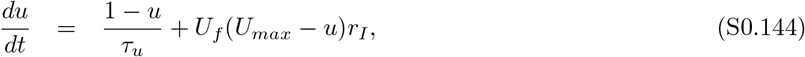

where *u* is the facilitation variable constrained to the interval [1, *U*_*max*_), *U*_*max*_ is the maximal facilitation value, *τ*_*u*_ is the time constant of STF, and *U*_*f*_ is the facilitation factor.

The Jacobian 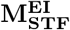 of the system with I-to-E STF is given by

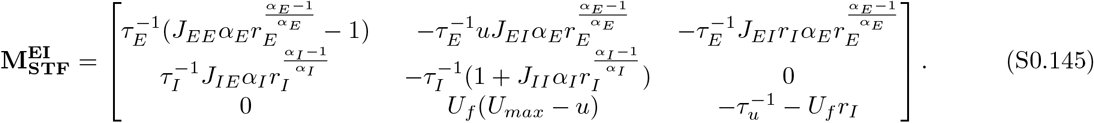

If inhibition is frozen, in other words, if feedback inhibition is absent, the Jacobian of the system becomes as follows:

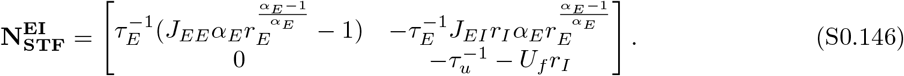

For the system with frozen inhibition, the dynamics are stable if

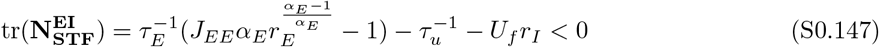

and

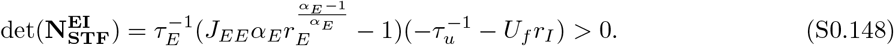

Therefore, if the network is an IS at the fixed point, the following condition has to be satisfied:

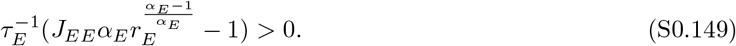

We further define the IS index for the system with I-to-E STF as follows:

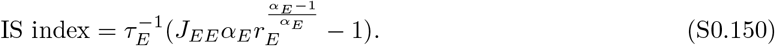

### Conditions for paradoxical response in networks with I-to-E STF

Next, we derived the condition of having the paradoxical effect in networks with I-to-E STF. Under the assumption that sufficient small perturbations would not lead to unstable network dynamics and regime transition between non-IS and IS, the conditions of having paradoxical effects are a positive slope of the excitatory nullcline and a larger slope of the inhibitory nullcline than that of the excitatory nullcline locally around the fixed point [17]. We can write the excitatory nullcline as follows

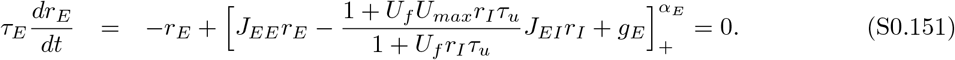

For *r*_*E*,*I*_ *>* 0, we have

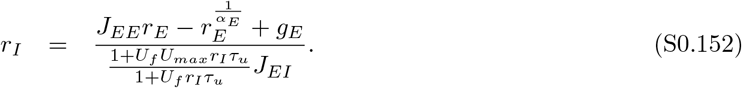

The slope of the excitatory nullcline in the *r*_*E*_*/r*_*I*_ plane where *x* axis is *r*_*E*_ and *y* axis is *r*_*I*_ can be written as follows

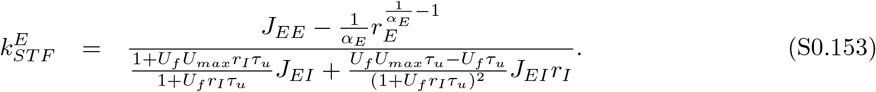

To have paradoxical effect, the slope of the excitatory nullcline at the fixed point of the system has to be positive. Therefore, we have

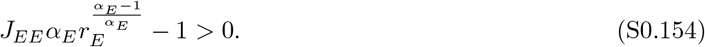

The inhibitory nullcline can be written as follows

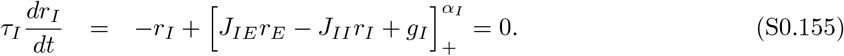

In the region of rates *r*_*E*,*I*_ *>* 0, we have

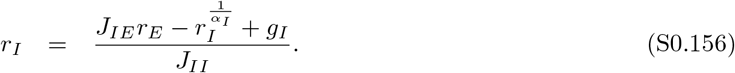

The slope of the inhibitory nullcline can be written as follows

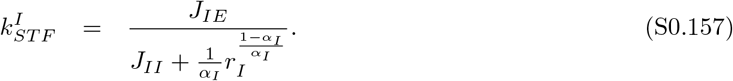

In addition to the positive slope of the excitatory nullcline, the slope of the inhibitory nullcline at the fixed point of the system has to be larger than the slope of the excitatory nullcline. We therefore have

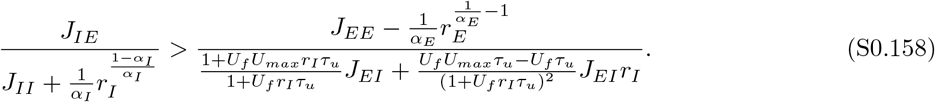

Note that to ensure that the system with I-to-E STF is stable, 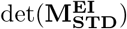 has to be negative. Therefore, we have

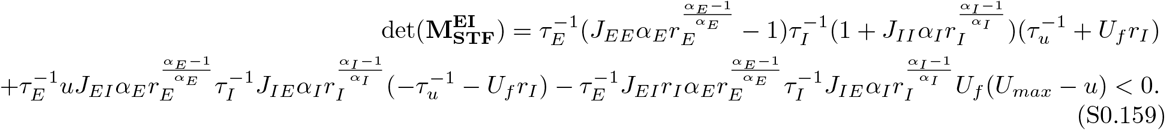

The condition shown in Eq. S0.158 is the same as the stability condition of the determinant of the Jacobian of the system with I-to-E STF, namely, as 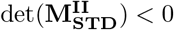. Thus, the condition is always satisfied when the system with I-to-E STF is stable.

The condition of being IS shown in Eq. S0.149 is identical to the condition of having paradoxical effect shown in Eq. S0.154, we therefore can conclude that in networks with I-to-E STF, inhibitory stabilization and the paradoxical effect imply each other.

### Conditions for IS in networks with I-to-I STD

The dynamics of networks with I-to-I STD are given by

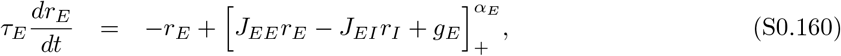

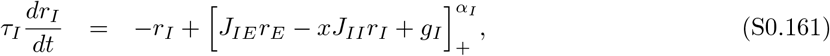

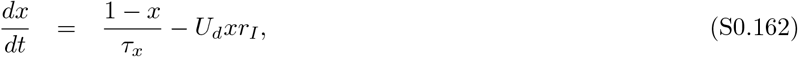

where *x* is the depression variable, which is limited to the interval (0, 1], *τ*_*x*_ is the depression time constant, and *U*_*d*_ is the depression factor.

The Jacobian 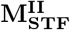 of the system with I-to-I STD is given by

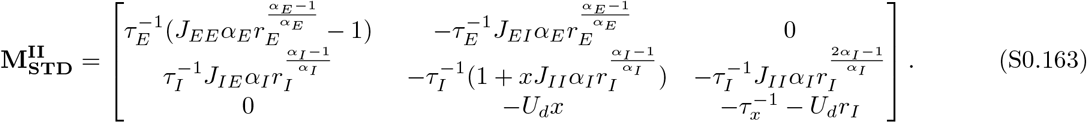

If inhibition is frozen, in other words, if feedback inhibition is absent, the Jacobian of the system becomes as follows:

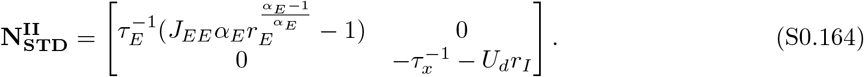

For the system with frozen inhibition, the dynamics are stable if

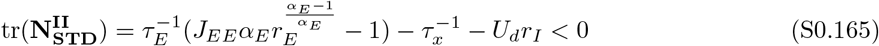

and

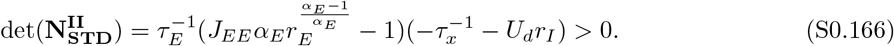

Therefore, if the network is an IS at the fixed point, the following condition has to be satisfied:

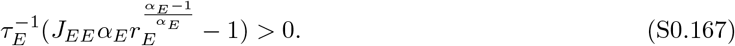

We further define the IS index for the system with I-to-I STD as follows:

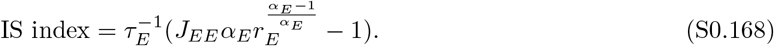

### Conditions for paradoxical response in networks with I-to-I STD

Next, we derived the condition of having the paradoxical effect in networks with I-to-I STD. Under the assumption that sufficient small perturbations would not lead to unstable network dynamics and regime transition between non-IS and IS, the conditions of having paradoxical effects are a positive slope of the excitatory nullcline and a larger slope of the inhibitory nullcline than that of the excitatory nullcline locally around the fixed point [17]. We can write the excitatory nullcline as follows

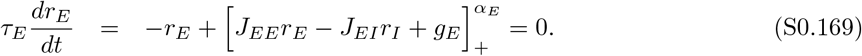

For *r*_*E*,*I*_ *>* 0, we have

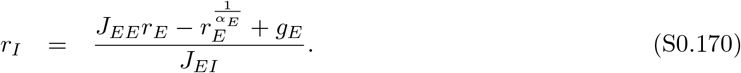

The slope of the excitatory nullcline in the *r*_*E*_*/r*_*I*_ plane where *x* axis is *r*_*E*_ and *y* axis is *r*_*I*_ can be written as follows

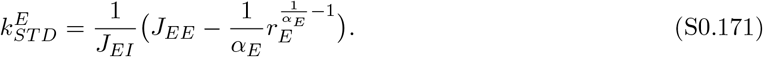

To have paradoxical effect, the slope of the excitatory nullcline at the fixed point of the system has to be positive. Therefore, we have

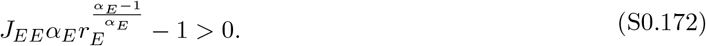

The inhibitory nullcline can be written as follows

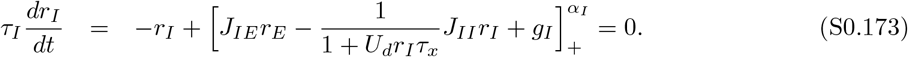

In the region of rates *r*_*E*,*I*_ *>* 0, we have

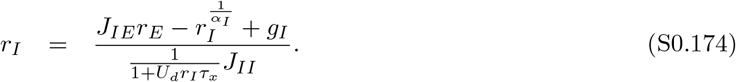

The slope of the inhibitory nullcline can be written as follows

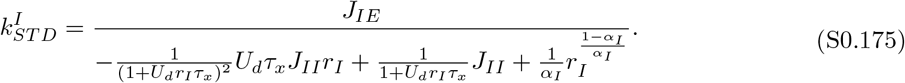

In addition to the positive slope of the excitatory nullcline, the slope of the inhibitory nullcline at the fixed point of the system has to be larger than the slope of the excitatory nullcline. We therefore have

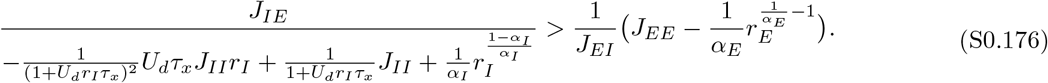

Note that to ensure that the system with I-to-I STD is stable, 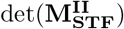 has to be negative. Therefore, we have

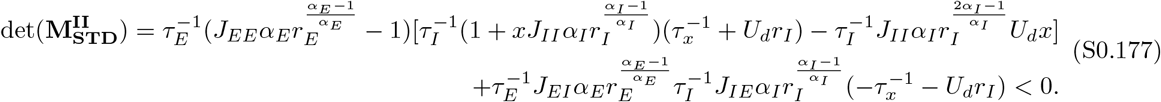

The condition shown in Eq. S0.176 is the same as the stability condition of the determinant of the Jacobian of the system with I-to-I STD, namely, as 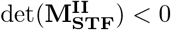. Thus, the condition is always satisfied when the system with I-to-I STD is stable.

The condition of being IS shown in Eq. S0.167 is identical to the condition of having paradoxical effect shown in Eq. S0.172, we therefore can conclude that in networks with I-to-I STD, inhibitory stabilization and the paradoxical effect imply each other.

### Conditions for IS in networks with I-to-I STF

The dynamics of networks with I-to-I STF are given by

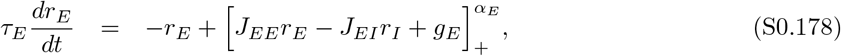

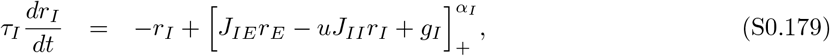

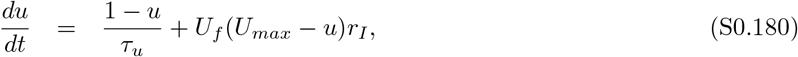

where *u* is the facilitation variable constrained to the interval [1, *U*_*max*_), *U*_*max*_ is the maximal facilitation value, *τ*_*u*_ is the time constant of STF, and *U*_*f*_ is the facilitation factor.

The Jacobian 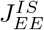 of the system with I-to-I STF is given by

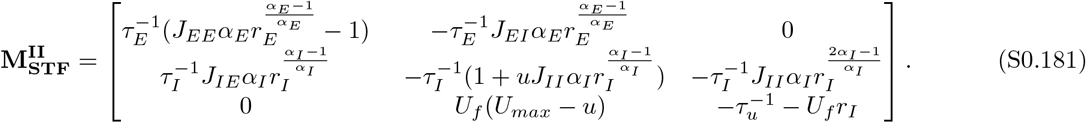

If inhibition is frozen, in other words, if feedback inhibition is absent, the Jacobian of the system becomes as follows:

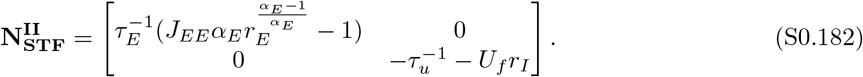

For the system with frozen inhibition, the dynamics are stable if

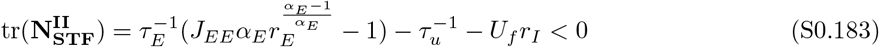

and

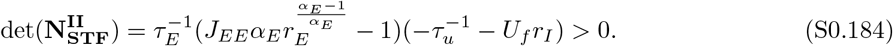

Therefore, if the network is an IS at the fixed point, the following condition has to be satisfied:

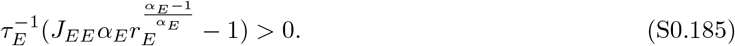

We further define the IS index for the system with I-to-I STF as follows:

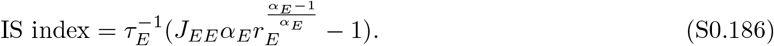

### Conditions for paradoxical response in networks with I-to-I STF

Next, we derived the condition of having the paradoxical effect in networks with I-to-I STF. Under the assumption that sufficient small perturbations would not lead to unstable network dynamics and regime transition between non-IS and IS, the conditions of having paradoxical effects are a positive slope of the excitatory nullcline and a larger slope of the inhibitory nullcline than that of the excitatory nullcline locally around the fixed point [17]. We can write the excitatory nullcline as follows

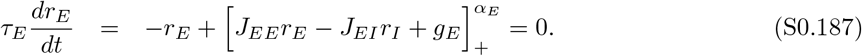

For *r*_*E*,*I*_ *>* 0, we have

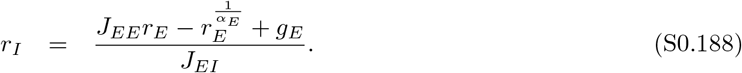

The slope of the excitatory nullcline in the *r*_*E*_*/r*_*I*_ plane where *x* axis is *r*_*E*_ and *y* axis is *r*_*I*_ can be written as follows

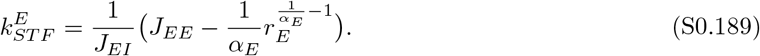

To have paradoxical effect, the slope of the excitatory nullcline at the fixed point of the system has to be positive. Therefore, we have

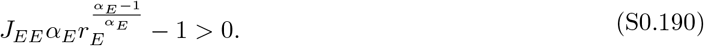

The inhibitory nullcline can be written as follows

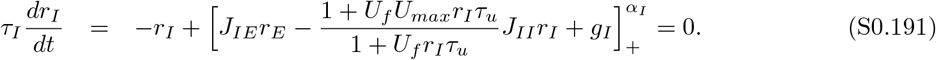

In the region of rates *r*_*E*,*I*_ *>* 0, we have

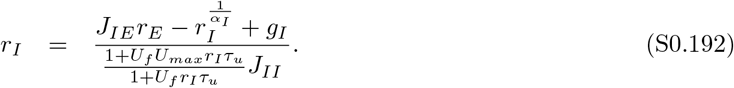

The slope of the inhibitory nullcline can be written as follows

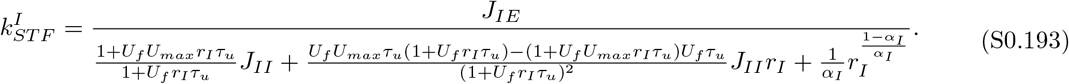

In addition to the positive slope of the excitatory nullcline, the slope of the inhibitory nullcline at the fixed point of the system has to be larger than the slope of the excitatory nullcline. We therefore have

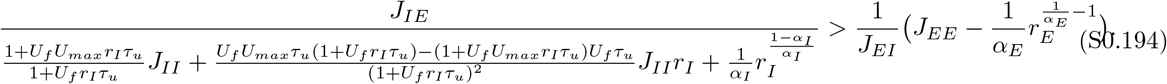

Note that to ensure that the system with I-to-I STF is stable, 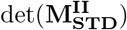 has to be negative. Therefore, we Have

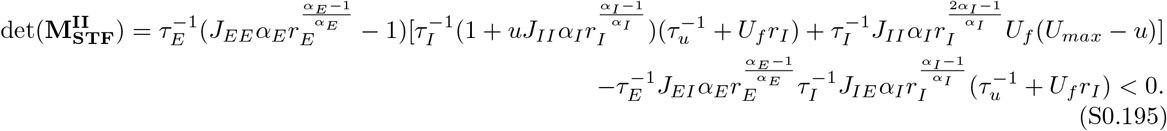

The condition shown in Eq. S0.194 is the same as the stability condition of the determinant of the Jacobian of the system with I-to-I STF, namely, as 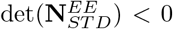. Thus, the condition is always satisfied when the system with I-to-I STF is stable.

The condition of being IS shown in Eq. S0.185 is identical to the condition of having paradoxical effect shown in Eq. S0.190, we therefore can conclude that in networks with I-to-I STF, inhibitory stabilization and the paradoxical effect imply each other.

### IS boundary for networks with E-to-I STD/STF, I-to-E STD/STF, and I-to-I STD/STF

We next investigated how the boundary between non-IS and IS, which we called ‘IS boundary’, changes as a function of *r*_*E*_. Mathematically, the IS boundary is determined by the corresponding recurrent excitatory-to-excitatory connection strength denoted by 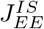 for different *r*_*E*_ at which the IS index is 0. Therefore, we have

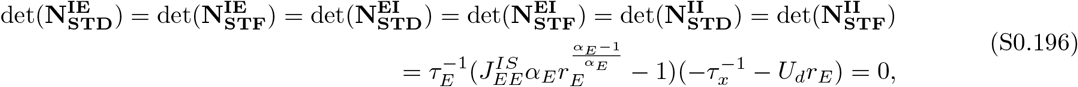

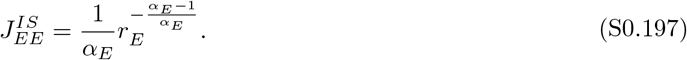

For *α* = 1, we have

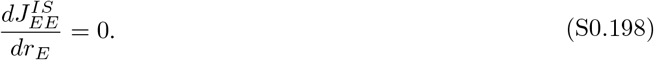

Therefore, for linear networks with E-to-I STD/STF, I-to-E STD/STF, and I-to-I STD/STF, the IS boundary does not change with *r*_*E*_.

For *α >* 1, we have

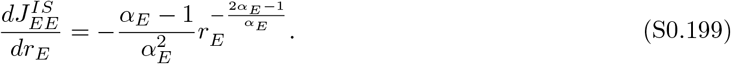

As a result, for supralinear networks with E-to-I STD/STF, I-to-E STD/STF, and I-to-I STD/STF, 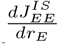 is always negative, suggesting that the boundary shifts downwards as *r*_*E*_ increases. Furthermore, the decrease of 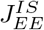 slows down for larger *r*_*E*_.

For *α <* 1, we have

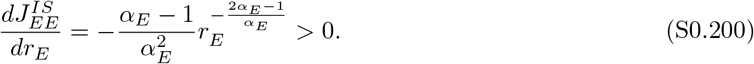

Therefore, for sublinear networks with E-to-I STD/STF, I-to-E STD/STF, and I-to-I STD/STF, the IS boundary always shifts upwards as *r*_*E*_ increases.

### Paradoxical boundary for networks with E-to-E STF, E-to-I STD/STF, I-to-E STD/STF, and I-to-I STD/STF

As inhibition stabilization and paradoxical effect imply each other in networks with E-to-E STF, E-to-I STD/STF, I-to-E STD/STF, and I-to-I STD/STF, the change of paradoxical boundary is identical to the change of IS boundary as we demonstrated before.

Note that conditions for IS and the paradoxical effect in networks with E-to-E STD and networks with E-to-I STF are shown in recent studies [29]. For the sake of completeness, we also included them in the Methods section.

### Generalization to networks with a mixture of short-term plasticity mechanisms

To generalize our results to networks with a mixture of short-term plasticity mechanisms, we described the dynamics of the network as follows:

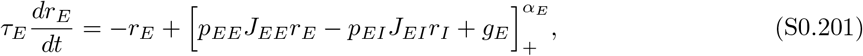

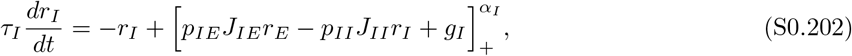

where *p*_*XY*_ denotes the short-term plasticity variable for the synapse from *Y* to *X*, with *X, Y ∈ {E, I}*. A *p*_*XY*_ larger than 1 suggests that the synapse is facilitating, whereas a *p*_*XY*_ less than 1 indicates that the synapse is depressing.

Here, we derived the condition of having the paradoxical effect. Under the assumption that sufficient small perturbations would not lead to unstable network dynamics and regime transition between non-IS and IS, the conditions of having paradoxical effects are a positive slope of the excitatory nullcline and a larger slope of the inhibitory nullcline than that of the excitatory nullcline locally around the fixed point. We can write the excitatory nullcline as follows

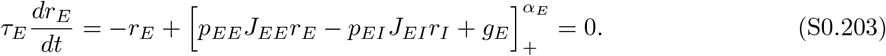

For *r*_*E*,*I*_ *>* 0, we have

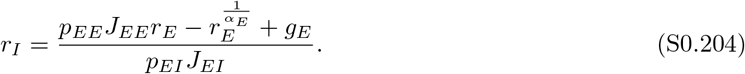

The slope of the excitatory nullcline in the *r*_*E*_*/r*_*I*_ plane where *x* axis is *r*_*E*_ and *y* axis is *r*_*I*_ can be written as follows

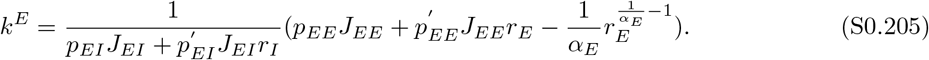

Provided that the inhibitory current to the excitatory network is always an increasing function (at least locally around the fixed point) of the inhibitory firing rate, we have

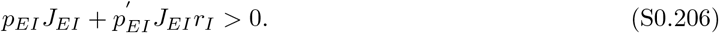

In addition to the positive slope of the excitatory nullcline, the slope of the inhibitory nullcline at the fixed point of the system has to be larger than the slope of the excitatory nullcline. This condition is satisfied as long as the stability of the network is granted.

Next, we sought to identify the relationship between the inhibition stabilization property of the network and the paradoxical effect.

In the presence of E-to-E STD and STP at other types of synapses, the condition of the paradoxical effect is

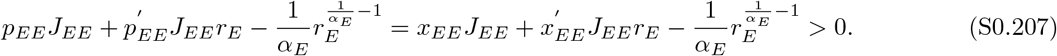

The above condition is the same as the condition that 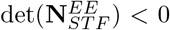. In networks with E-to-E STD as well as STP at other types of synapses, the effect of STP at other types of synapses does not act on the excitatory subnetwork when feedback inhibition is impermissible. Thus, the IS conditions for networks with E-to-E STD and STP at other types of synapses are the same as these for networks with E-to-E STD alone. As we showed before, 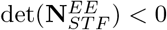 is the sufficient condition for IS in networks with E-to-E STD, but not the necessary condition. Therefore, in the presence of E-to-E STD and STP at other types of synapses, the paradoxical effect implies inhibition stabilization, whereas inhibition stabilization does not necessarily imply the paradoxical effect.

In the presence of E-to-E STF and STP at other types of synapses, the condition of the paradoxical effect changes from Eq. S0.207 to

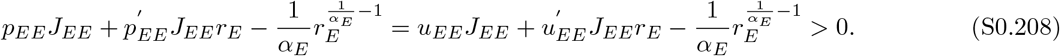

The above condition is the same as the condition that 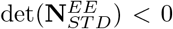. In networks with E-to-E STF and STP at other types of synapses, the effect of STP at other types of synapses does not act on the excitatory subnetwork when feedback inhibition is impermissible. Thus, the IS conditions for networks with E-to-E STF and STP at other types of synapses are the same as these for networks with E-to-E STF alone. As we showed before, 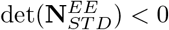 is the necessary and sufficient condition for IS in networks with E-to-E STF. Therefore, in the presence of E-to-E STF and STP at other types of synapses, inhibition stabilization and the paradoxical effect imply each other.

Similarly, in the absence of E-to-E STP but in the presence of STP at other types of types of synapses, the condition of the paradoxical effect instead becomes

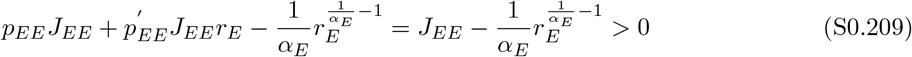

In networks without E-to-E STD but with STP at other types of synapses, the effect of STP at other types of synapses does not act on the excitatory subnetwork when feedback inhibition is impermissible. The IS conditions for networks without E-to-E STP but with STP at other types of synapses are the same as these for networks with static connectivity. The above condition is the necessary and sufficient condition for IS in networks with static connectivity. Therefore, in the absence of E-to-E STP but in the presence of STP at other types of synapses, inhibition stabilization and the paradoxical effect imply each other.

### Generalization to networks with both E-to-E STD and E-to-E STF

To generalize our results to networks with both STD and STF on the E-to-E synapses, we described the dynamics of the network as follows:

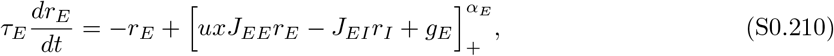

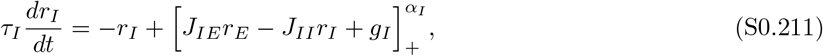

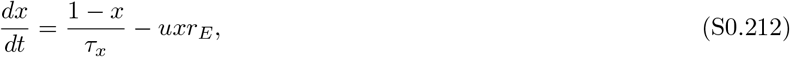

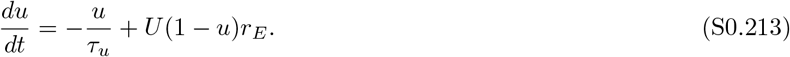

The Jacobian 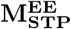 of the system with E-to-E STP is given by

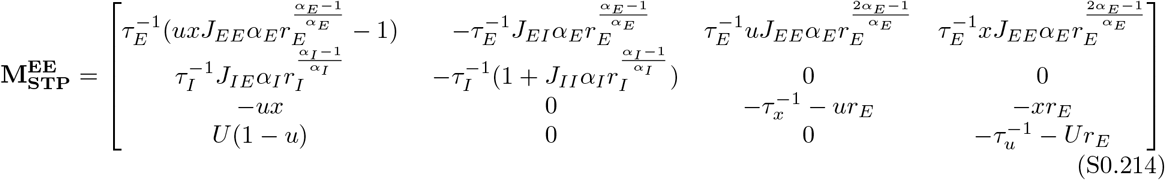

If inhibition is frozen, i.e., if feedback inhibition is absent, the Jacobian of the system becomes as follows:

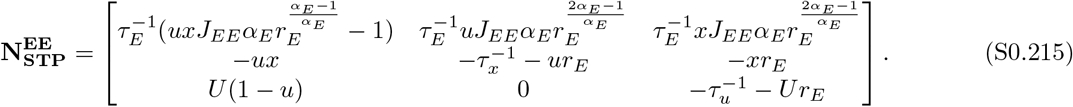

As the product of three eigenvalues of 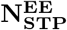 is the determinant 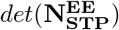, the network is inhibition stabilized if 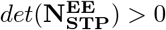 so we can write:

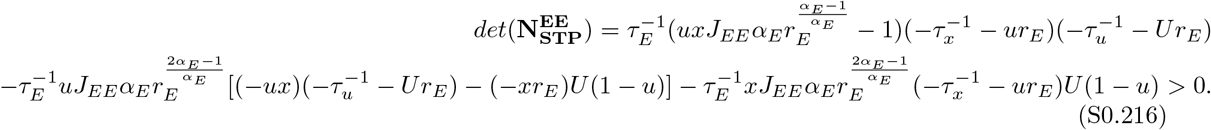

Therefore, we have

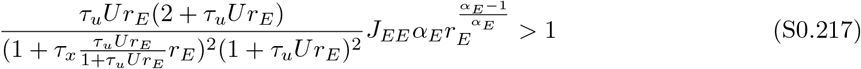

As the sum of three eigenvalues of 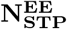 is the trace 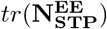, the network is inhibition stabilized if 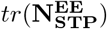 and we can write:

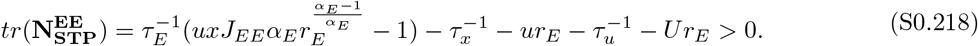

Therefore, we have

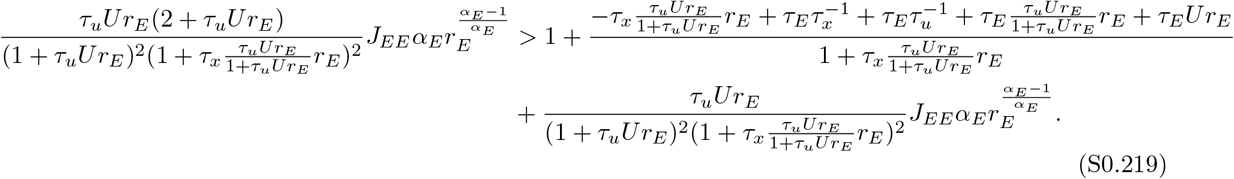

Note that the trace condition is independent from the determinant condition. For instance, in the limit of small *τ*_*x*_ and large *τ*_*u*_ when E-to-E STD dominates over E-to-E STF, we have:

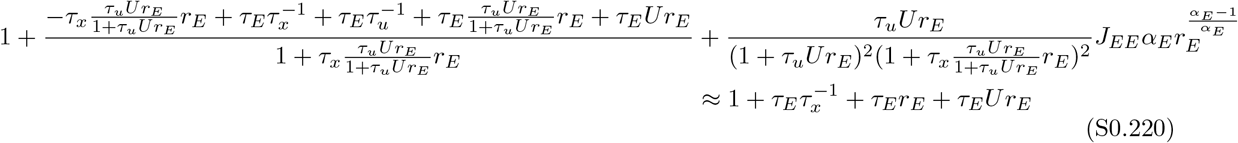

The trace condition then becomes as follows

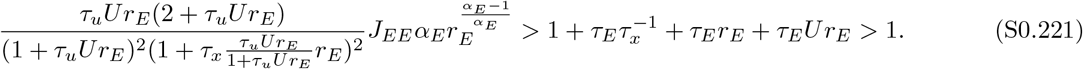

Therefore, when E-to-E STD dominates over E-to-E STF, the network can be inhibition stabilized by satisfying the determinant condition, but it does not have to satisfy the trace condition.

Furthermore, it is worth mentioning that as 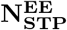 is a 3 by 3 matrix, the determinant and trace conditions are sufficient conditions to identify IS but not the necessary conditions.

Next, we derived the condition of having the paradoxical effect. Under the assumption that sufficient small perturbations would not lead to unstable network dynamics and regime transition between non-IS and IS, the conditions of having paradoxical effects are a positive slope of the excitatory nullcline and a larger slope of the inhibitory nullcline than that of the excitatory nullcline locally around the fixed point. We can write the excitatory nullcline as follows

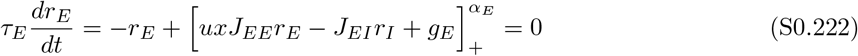

For *r*_*E*,*I*_ *>* 0, we have

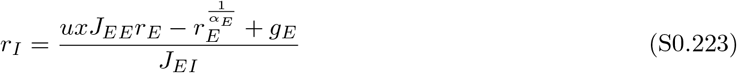

The slope of the excitatory nullcline in the *r*_*E*_*/r*_*I*_ plane where *x* axis is *r*_*E*_ and *y* axis is *r*_*I*_ can be written as follows

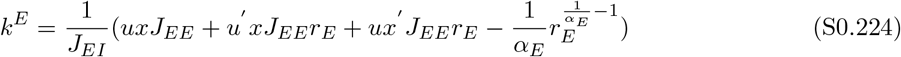

To have paradoxical effect, the slope of the excitatory nullcline at the fixed point of the system has to be positive. Therefore, we have

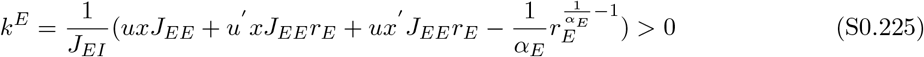

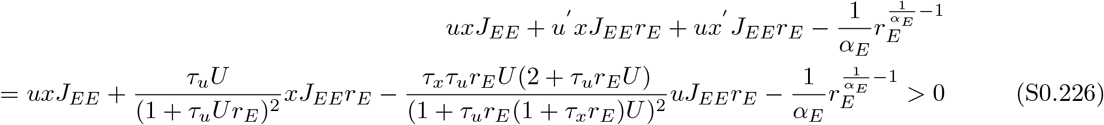

We can then obtain the condition below:

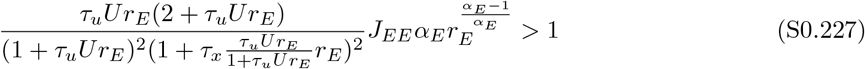

This condition is identical to the determinant condition for being IS.

In addition to the positive slope of the excitatory nullcline, the slope of the inhibitory nullcline at the fixed point of the system has to be larger than the slope of the excitatory nullcline. This condition is satisfied as long as the stability of the network is granted.

Therefore, in the presence of E-to-E STD and STF, the paradoxical effect implies inhibition stabilization, whereas inhibition stabilization does not necessarily imply the paradoxical effect.

### Generalization to networks with multiple inhibitory cell types

To generalize our results to networks with multiple inhibitory cell types, we considered a network with one excitatory population and *N −*1 inhibitory populations. The dynamics of the network can be described as follows:

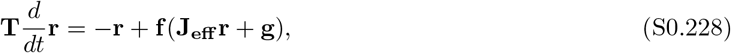

where **T** is a diagonal matrix containing the time constants of firing rate dynamics for different populations, **r** is a vector containing the firing rates of different populations, **f** (**x**) is a vector containing the input-output functions for different populations, **g** is a vector containing the inputs to different populations.

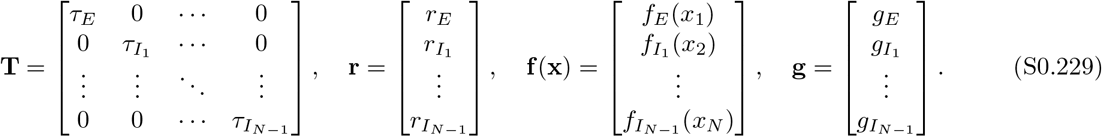

**J**_**eff**_ is the effective connectivity matrix defined as follows:

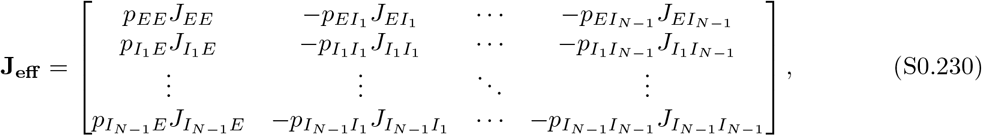

where *p*_*XY*_ denotes the short-term plasticity variable for the synapse from *Y* to *X*, with *X, Y ∈ {E, I*_1_, *· · ·, I*_*N−*1_*}*. A *p*_*XY*_ larger than 1 suggests that the synapse is facilitating, whereas a *p*_*XY*_ less than 1 indicates that the synapse is depressing.

The equation linearized about the fixed point can be written as follows:

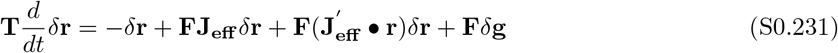

where *δ***r** is a vector containing the changes in firing rates of different populations, *δ***g** is a vector containing the changes in the inputs to different populations, **F** is a diagonal matrix containing the derivatives of the input-output functions at the fixed point:

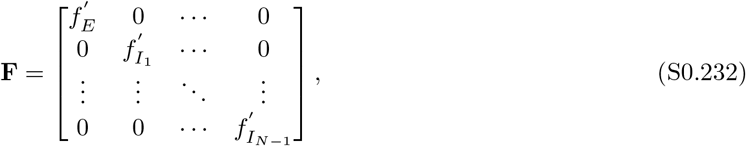

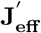 is a tensor and given by

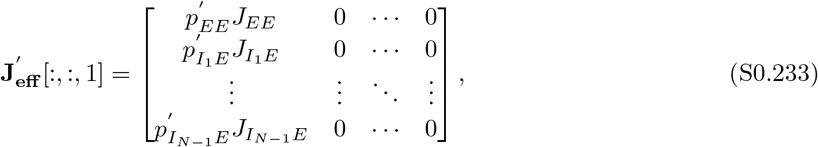

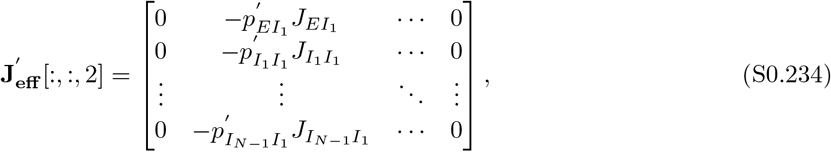

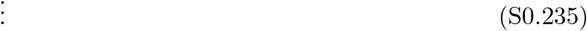

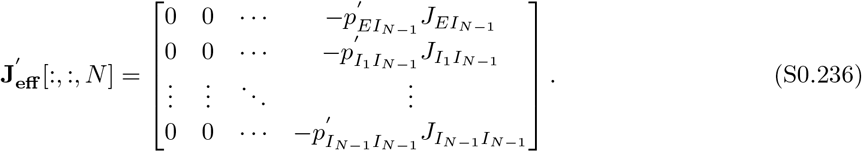

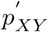 denotes the derivative of the short-term plasticity variable *p*_*XY*_ with respect to *r*_*Y*_ . 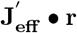 is a 3-mode product of the tensor 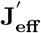 and the vector **r**:

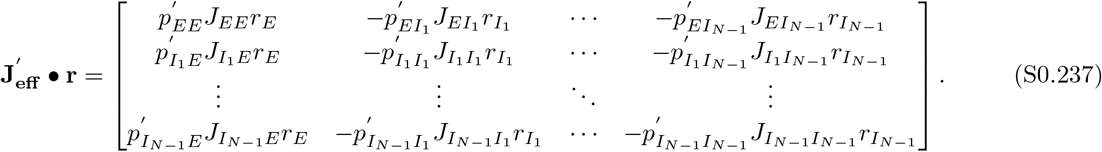

If only inhibitory populations receive external perturbative inputs, i.e., *δg*_*E*_ is 0, we have:

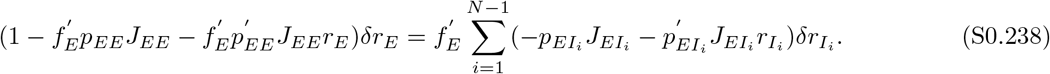

Note that 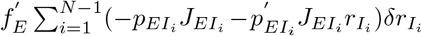 represents the change in the total inhibitory current to the excitatory population. If it is negative, the total inhibitory current to the excitatory population increases.

Provided that 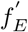 is positive, namely, the input-output function is a monotonically increasing function, we have the following two cases:

1. If 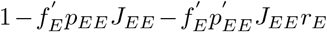 is negative, then the sign of the change in the firing rate of the excitatory population and of the change in the total inhibitory current to the excitatory population are the same. This implies that if 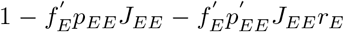 is negative, additional inputs to the inhibitory populations increase or decrease the firing rate of the excitatory population as well as the total inhibitory current to the excitatory population.
2. If 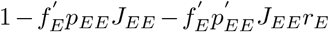 is positive, then the sign of the change in the firing rate of the excitatory population and of the change in the total inhibitory current to the excitatory population are opposite.

Next, we sought to identify the relationship between the sign of 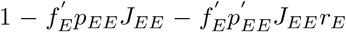 and the inhibition stabilization property of the network.

In the presence of E-to-E STD and STP at other types of synapses, we have

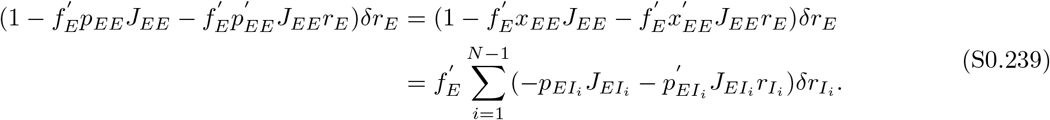

In this case, the condition that 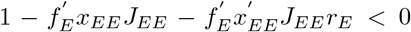 is the same as the condition that 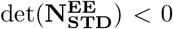. In networks with E-to-E STD and STP at other types of synapses, the effect of STP at other types of synapses does not act on the excitatory subnetwork when feedback inhibition is not allowed (frozen inhibition). Thus, the IS conditions for networks with E-to-E STD and STP at other types of synapses are the same as these for networks with E-to-E STD alone. As we showed before, 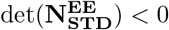 is a sufficient condition for IS in networks with E-to-E STD, but not a necessary condition. Therefore, in the presence of E-to-E STD, if the network is IS, the sign of the change in the firing rate of the excitatory population and of the change in the total inhibitory current to the excitatory population are the same. However, the same sign of the change in the firing rate of the excitatory population and of the change in the total inhibitory current to the excitatory population does not imply that the network is IS.

In the presence of E-to-E STF and STP at other types of synapses, we have:

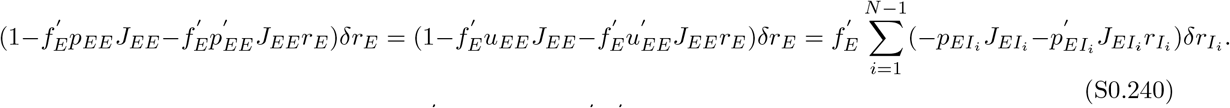

In this case, the condition that 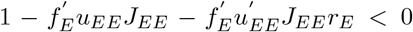 is the same as the condition that 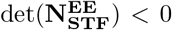. In networks with E-to-E STF and STP at other types of synapses, the effect of STP at other types of synapses does not act on the excitatory subnetwork when feedback inhibition is not allowed. Thus, the IS conditions for networks with E-to-E STF and STP at other types of synapses are the same as those for networks with E-to-E STF alone. As we showed before, 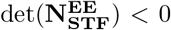 is a necessary and a sufficient condition for IS in networks with E-to-E STF. Therefore, in the presence of E-to-E STF, if the network is non-IS, then the sign of the change in the firing rate of the excitatory population and of the change in the total inhibitory current to the excitatory population are opposite. And if the network is IS, the sign of the change in the firing rate of the excitatory population and of the change in the total inhibitory current to the excitatory population are the same.

In the absence of E-to-E STP but in the presence of STP at other types of synapses, we have:

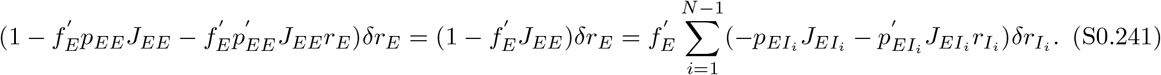

In networks without E-to-E STD but with STP at other types of synapses, the effect of STP at other types of synapses does not act on the excitatory subnetwork when feedback inhibition is impermissible. The IS conditions for networks without E-to-E STP but with STP at other types of synapses are the same as these for networks with static connectivity. The above condition is a necessary and a sufficient condition for IS in networks with static connectivity. Therefore, in the absence of E-to-E STP but in the presence of STP at other types of synapses, if the network is non-IS, then the sign of the change in the firing rate of the excitatory population and of the change in the total inhibitory current to the excitatory population are opposite. And if the network is IS, the sign of the change in the firing rate of the excitatory population and of the change in the total inhibitory current to the excitatory population are the same.

## Notes

### Competing Interest Statement

The authors have declared no competing interest.

### Summary of Updates

Added generalizations to more complex scenarios.

